# Epithelial colonies *in vitro* elongate through collective effects

**DOI:** 10.1101/755181

**Authors:** Jordi Comelles, SS Soumya, Linjie Lu, Emilie Le Maout, S. Anvitha, Guillaume Salbreux, Frank Jülicher, Mandar M. Inamdar, Daniel Riveline

## Abstract

Epithelial tissues of the developing embryos elongate by different mechanisms, such as neighbor exchange, cell elongation, and oriented cell division. Since autonomous tissue self-organization is influenced by external cues such as morphogen gradients or neighboring tissues, it is difficult to distinguish intrinsic from directed tissue behavior. The mesoscopic processes leading to the different mechanisms remain elusive. Here, we study the spontaneous elongation behavior of spreading circular epithelial colonies in vitro. By quantifying deformation kinematics at multiple scales, we report that global elongation happens primarily due to cell elongations, and its direction correlates with the anisotropy of the average cell elongation. By imposing an external time-periodic stretch, the axis of this global symmetry breaking can be modified and elongation occurs primarily due to orientated neighbor exchange. These different behaviors are confirmed using a vertex model for collective cell behavior, providing a framework for understanding autonomous tissue elongation and its origins.

## Introduction

Tissue elongation is a central morphogenetic event occurring in many organisms during development (Lecuit and Lenne, 2007; Guillot and Lecuit, 2013), such as *Drosophila* or *C. elegans*. The tissue is transformed both in terms of area and shape. Such transformation takes place within typically hour timescale with or without cell division. During this process, symmetry of cells and tissues is broken by different mechanisms, such as neighbor exchange (Rauzi *et al.*, 2008; Rauzi, Lenne and Lecuit, 2010), cell elongation (Clarletta, Ben Amar and Labouesse, 2009; Vuong-Brender, Yang and Labouesse, 2016) and oriented cell division (Campinho *et al.*, 2013). Rearrangement of neighboring cells or T1 transitions is essential in the germ band extension of *Drosophila* (Rauzi *et al.*, 2008; Rauzi, Lenne and Lecuit, 2010), allowing a group of cells to change their position by intercalation, eventually leading to tissue elongation. Cell deformation drives the 3-fold elongation process in *C. elegans* (Clarletta, Ben Amar and Labouesse, 2009; Vuong-Brender, Yang and Labouesse, 2016) while keeping the number of cells and their positions fixed. Finally, epithelial spreading during zebrafish epiboly is promoted by oriented cell divisions as a mechanism to limit tension (Campinho *et al.*, 2013). Those mechanisms can act alone or in combination as in *Drosophila* pupal wing elongation (Etournay *et al.*, 2015). While the phenomenon is known to involve remodeling of adherens junctions (Rauzi, Lenne and Lecuit, 2010) and acto-myosin (He *et al.*, 2010; Rauzi, Lenne and Lecuit, 2010) at the molecular level, mesoscopic mechanisms leading to distinct morphogenesis processes are poorly understood. This is partly because inputs from morphogen gradients (Gilmour, Rembold and Leptin, 2017) or from neighboring tissues (Zhang *et al.*, 2011; Etournay *et al.*, 2015) can affect tissue autonomous self-organization *in vivo*. For example, changes in tissue shape can be influenced by neighboring tissues such as the cuticle and the blade in the *Drosophila* pupal wing elongation (Etournay *et al.*, 2015), the coordination between amnioserosa and epidermis in dorsal closure (Hayes and Solon, 2017), and the muscle layer in gut development (Shyer *et al.*, 2013) or in *C. elegans* morphogenesis (Zhang *et al.*, 2011). Since *in vivo*, epithelial tissues are surrounded by other tissues and the cellular dynamics leading to elongation can result from interactions between tissues and boundary conditions, it is therefore difficult to disentangle cell intrinsic from externally mediated behaviors. In this context, it appears important to characterize elongation in an *in vitro* system where the epithelial tissue undergoes shape transition autonomously.

Here we use *in vitro* experiments and numerical simulations to characterize the spontaneous behavior of a growing cell colony *in vitro*. We designed an assay to study the spontaneous elongation of a tissue that is not subjected to external orienting input, we studied the appearance of the symmetry breaking, and the effect that external forces have in this process. We show that *in vitro* tissue elongation arises from anisotropy in the average cell elongation. This anisotropy sets the direction along which boundary cells migrate radially outwards resulting in a non-isotropic elongation that arises primarily through cell elongation. For colonies submitted to a time periodic uniaxial stretch, the axis of global symmetry breaking can be imposed by external force, and tissue elongation arises through oriented neighbor exchange. Emergence of radially migrating cells and the interplay between cell elongation and cell rearrangements are confirmed by numerical simulations based on a vertex model. Our results suggest that spontaneous shape deformation is related to the mean orientation of the nematic cell elongation field in the absence of any external input. This provides a framework to explain autonomous tissue elongation and how contributions from different mesoscopic mechanisms can be modulated by external forces.

## Results

### Isotropic colonies elongate in a non-isotropic manner

To study the spontaneous tissue deformation arising during epithelial growth, we designed an *in vitro* assay to track symmetry breaking, both spontaneous and driven by external force. We prepared isotropic colonies of Madin Darby Canine Kidney (MDCK) cells, which assume features of epithelial cells *in vivo* (Reinsch and Karsenti, 1994; Adams *et al.*, 1998; Reffay *et al.*, 2014), such as adherens junctions (Adams *et al.*, 1998), cytoskeletal components and the Rho signaling pathway regulating cell shapes and dynamics (Reffay *et al.*, 2014; Fodor *et al.*, 2018). The initial size and shape of the colonies were controlled by plating cells in microfabricated circular stencils (Ostuni *et al.*, 2000). When cells reached confluency, the stencil was removed at time *t*_0_. Cell dynamics was followed over time by phase contrast (Video 1) or fluorescence microcopy with strains labeled with GFP cadherin (Figure 1b), that allowed to observe the behavior of individual cells. We observed that large colonies (750 μm in diameter) expanded isotropically (Figure 1 – figure supplement 1). In contrast, colonies of 250 μm in diameter (Figure 1a), the typical coherence length of such epithelial tissues (Doxzen *et al.*, 2013), expanded in a non-isotropic manner (Figure 1c). We then further characterized the process of symmetry breaking.

**Figure 1.**
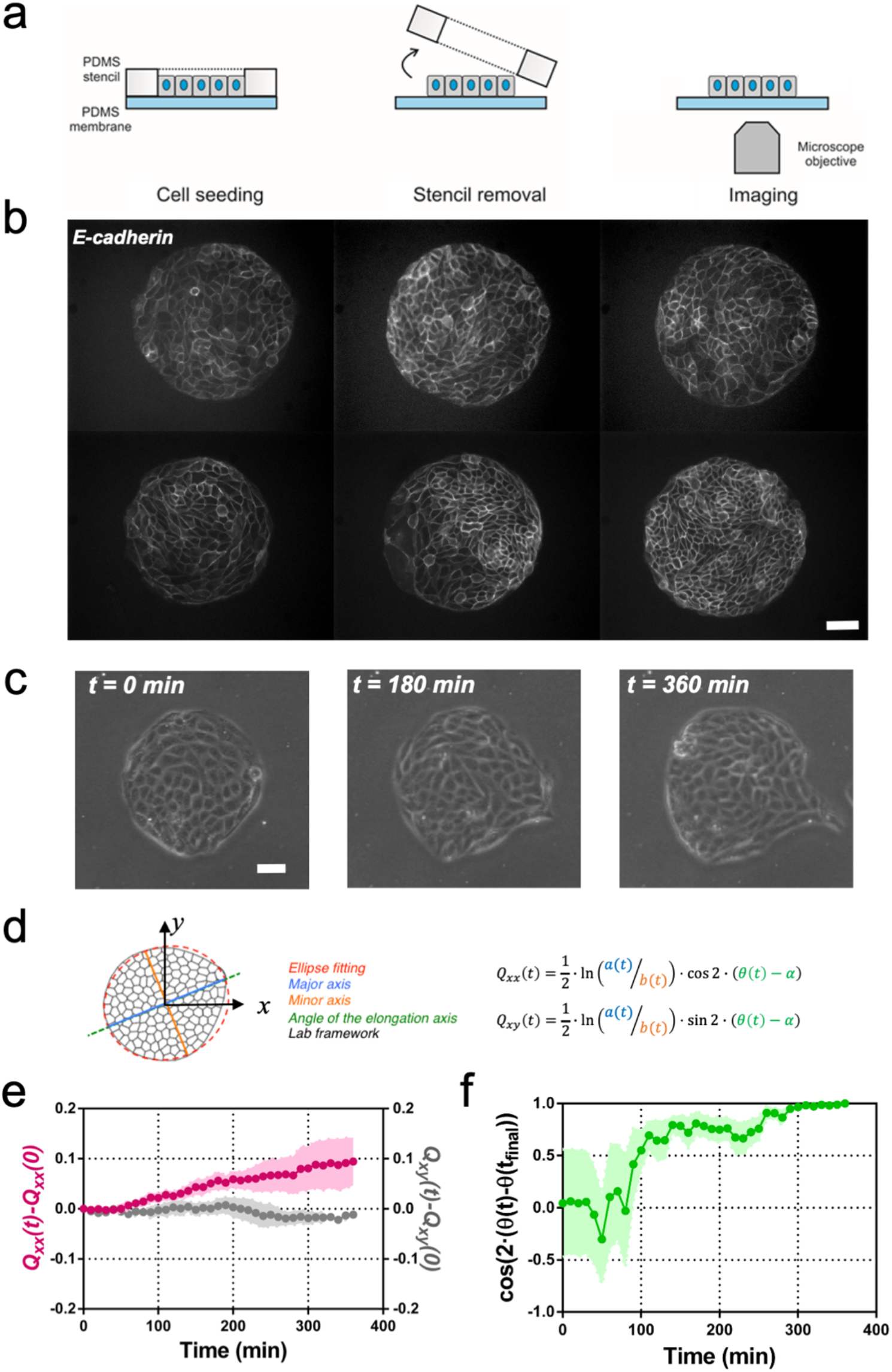
Symmetry breaking and its quantification. (**a**) Schematics of the experimental set-up: MDCK cells were seeded on a PDMS membrane using stencils to predefine their shape. When the colony was confluent, the stencil was removed and the expansion of the colony was observed under the microscope. (**b**) Several examples of MDCK colonies (GFP-E-cadherin) after stencil removal and prior to colony expansion. Scale bar 50 μm. (**c**) Phase contrast images of the spontaneous elongation of a MDCK colony for 360 min. Scale bar 50 μm. (**d**) Colony elongation is quantified by ellipse fitting and *Q_xx_* and *Q_xy_* measurement referred to the elongation axis (*α* = *θ* (t_final_)). (**e**) *Q_xx_* (left y axis) and *Q_xy_* (right y axis) during 360 min of colony expansion. Mean value ± standard error of the mean, n = 4 colonies from N = 4 independent experiments. (**f**) Cosine of two times the angle difference between the instantaneous main direction of the colony (*θ* (t)) and the main direction of the colony at 360 min (*θ* (t_final_)). Colonies set the elongation direction within the first 120 min. Mean value ± standard error of the mean, n = 4 colonies from N = 4 independent experiments.

In order to compare elongations in each experiment, we quantified the breaking of symmetry by ellipse-fitting the colony shape. Shape change analysis was quantified by a nematic shape elongation tensor ****Q****. It has two independent components defined as *Q_xx_* = ½ ln(*a/b*)cos(*2•(θ − α*) and *Q_xy_ =* ½ ln(*a/b*)sin (*2•(θ − α*), where *a* corresponds to the major axis, *b* to the minor axis, *θ* to the orientation of the major axis of the ellipse and *α* = *θ* (t_final_) (Figure 1d). As can be seen in Figure 1e, MDCK colonies elongated persistently along the main axis of elongation (*Q_xx_* > 0 and *Q_xy_* ≈ 0) for 6 hours (Figure 1e). In addition, we explored if other epithelial cell lines would behave in a similar manner. Circular epithelial colonies of human epithelial colorectal adenocarcinoma cells (Caco2) and human mammary epithelial cells (MCF-10A) also elongated along the main axis of elongation and by the same magnitude that MDCK cells (Figure 1 – figure supplement 2). We note that elongation observed during this time for the three epithelial cell lines was similar in magnitude to tissue elongation observed during *in vivo* morphogenesis, for instance in the wing blade in *Drosophila* (Etournay *et al.*, 2015). Moreover, the elongation direction (*θ*_final_ =*θ* (*t* = *6* h)) converges to a constant value within 2 hours after *t*_0_ (Figure 1f). Altogether, large circular epithelial colonies (750 μm in diameter) expand isotropically, whereas small colonies (250 μm in diameter) expand in a anisotropic manner and shape symmetry breaking takes place within the first 2 hours. As a result, we focus here on these first 2 hours during which the elongation axis is established.

### Cyclic uniaxial stretching rectifies symmetry breaking

It has been previously described for *C. elegans* embryo elongation (Zhang *et al.*, 2011) and in other organisms (Zhang and Labouesse, 2012) that time periodic stretch can play a role in morphogenesis. Motivated by these observations, we explored whether oscillatory external forces could have an impact on the direction of elongation. We designed an experimental set-up where elongating colonies were submitted to cyclic uniaxial stretching (Figure 2a and Video 2). Mechanical cycles of contraction-relaxation can range from 1 s in *C. elegans* epithelial elongation (Zhang *et al.*, 2011) up to 200 s in dorsal closure (Solon *et al.*, 2009). So, we explored frequencies and extensions around physiological values (Zhang and Labouesse, 2012). We selected three different cycle durations (20 s, 60 s, and 120 s period) and three different stretching conditions (5%, 10% and 15% strain). The stretch was applied to a silicon membrane and was transmitted to the colony. We then fitted the colonies shapes with ellipses at successive time and quantified *Q_xx_* and *Q_xy_* with respect to the angle of uniaxial stretching (set as *x*-axis, *α* = 0). Figure 2 – figure supplement 1 shows the value of the components of the tensor ****Q**** along time for the different strains and periods tested. Among different conditions, we observed colony elongation along the direction imposed by the external strain when we stretched cyclically with 60 s timescale and 5% strain (Figure 2b and Video 3). The overall elongation of colonies under cyclic uniaxial stretching was similar to the spontaneous elongation in the absence of externally applied uniaxial stretching during the first two hours (Figure 2c). Also, the magnitude of the shape elongation tensor ****Q**** under cyclic uniaxial stretching was comparable to the spontaneous elongation of colony when stretch was not applied, but elongation was oriented in the direction of externally applied uniaxial cyclic stretching (Figure 2d). Therefore, application of an external cyclic force can rectify symmetry breaking and set the direction of tissue elongation.

**Figure 2.**
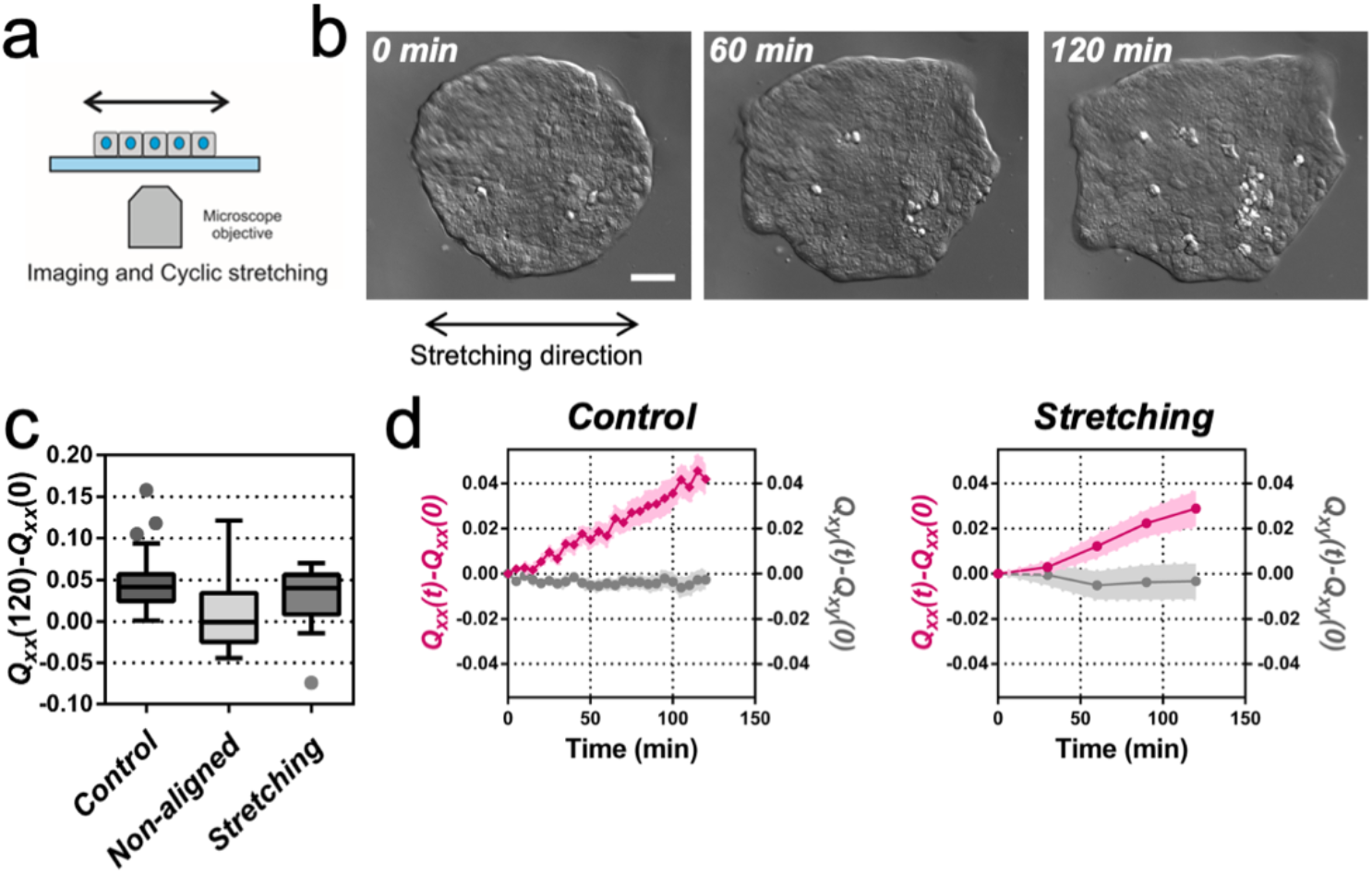
Uniaxial cyclic stretching rectifies symmetry breaking. (**a**) Schematics of the experiment where the colony expansion was observed under the microscope while the underlying membrane was uniaxially stretched. (**b**) Snapshots of the expansion of a MDCK colony while cyclically stretched. Scale bar 50 μm. (**c**) Colony elongation (*Q_xx_*) of control colonies along the elongation axis, control colonies in the laboratory framework (non-aligned, *α* = 0) and colonies under cyclic uniaxial stretching in the laboratory framework (uniaxial stretching, *α* = 0). Box Plot between 25th and 75th percentile, being the line in the box the median value, whiskers and outliers (dots) are obtained following Tukey’s method, N_control_ = 11 independent experiments, n = 25 colonies and N_stretching_ = 9, n = 20 colonies. Mann-Whitney test control *vs* control aligned p = 0.0003, control *vs* stretching p = 0.0281 and control aligned *vs* stretching p = 0.3319. (**d**) *Q_xx_* (left y axis) and *Q_xy_* (right y axis) during 120 min of colony expansion for control colonies and colonies under cyclic uniaxial stretching. Mean value ± standard error of the mean, N_control_ = 8, n = 14 colonies and N_stretching_ = 9, n = 20 colonies.

### Collective effects are essential for rectification

To get further insight into the collective nature of the rectification of tissue elongation, we probed the roles of adhesion between cells. First, we stretched single MDCK cells, individually plated. We observed that cells oriented perpendicularly to the externally applied uniaxial cyclic stretching (Figures 3a and 3b) as previously reported for fibroblasts (Faust *et al.*, 2011).Then we blocked cell-cell junction in circular colonies prior stretching. Briefly, we incubated confluent colonies in medium containing 5 mM EDTA and 10 μg/ml anti-E-cadherin blocking antibody that targeted the extracellular domain of E-cadherin for 30 min (Harris, Daeden and Charras, 2014). Then, medium was replaced by normal medium and the evolution of colonies with and without stretch was followed (Figure 3c). In the absence of externally applied uniaxial cyclic stretching, colonies treated with anti E-cadherin antibody expanded more than control colonies. Moreover, this expansion was still along one preferential direction (Figure 3d). Under cyclic uniaxial stretching, elongation was also non-isotropic and along the direction perpendicular to the cyclic stretching direction, in contrast to the parallel elongation observed when cell-cell contacts were intact (Figure 2b–d). This supports the collective nature of colony elongation. It is worth noting that cells inside the colony exhibited a decrease in their mean velocity (Figure 3e) and a large recruitment of myosin within cells similar to reinforcements (Riveline *et al.*, 2001) in stretching conditions, as shown by the appearance of stress fibers (Figure 3f). However, this effect did not appear to affect the overall elongation rate. Altogether these data suggest that the asymmetric expansion of colonies in the direction imposed by cyclic uniaxial stretching is generally associated to a collective effect.

**Figure 3.**
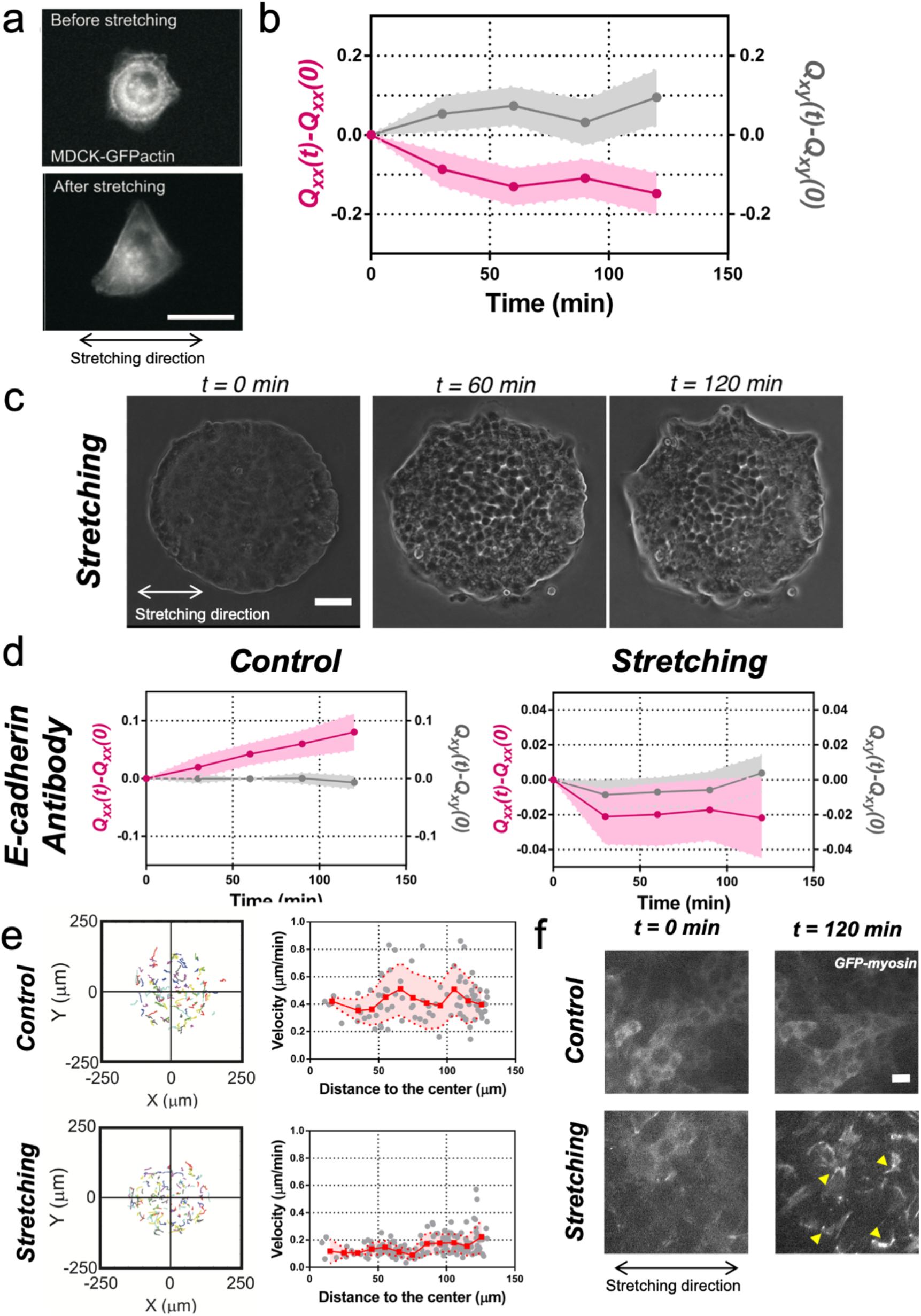
Collective effects are essential for rectification. (**a**) Image of a single MDCK-GFP-actin cell before and after being stretch for 2 hours (5% strain and 60s period). Scale bar 20μm. (**b**) *Q_xx_* (left y axis) and *Q_xy_* (right y axis) of single cells during 120 min of cyclic uniaxial stretching (*α* = 0). Mean value ± standard error of the mean. N = 3, n = 31 cells. (**c**) Phase contrast images of an MDCK colony evolving for 120 min under cyclic mechanical stretching when E-cadherins are blocked by an E-cadherin antibody. Scale bar 50μm. (**d**) Comparison of the cumulative *Q_xx_* (left y axis) and cumulative *Q_xy_* (right y axis) during 120 min of colony expansion when E-cadherin are blocked by an E-cadherin antibody in control and under cyclic uniaxial stretching (*α* = 0). Mean value ± standard error of the mean, N_control_ = 3, n = 8 colonies and N_stretching_ = 4, n = 15 colonies. (**e**) Trajectories of cells (left) and single cell velocity as a function of its distance to the center of the colony (right) in control colonies (top) and under cyclic uniaxial stretching (bottom). n_control_ = 90 cells from 8 colonies of N_control_ = 4 independent experiments and n_stretching_ = 154 cells from 13 colonies of N_stretching_ = 4 independent experiments. Individual cells in grey, red square and line corresponds mean (binned by distance to the center), shadowed area corresponds to SD. (**f**) Myosin distribution inside MDCK-GFP-myosin colonies at 0 min and at 120 min after expansion in control and under uniaxial stretching. Note the myosin structures appearing in the stretching case (yellow arrows). Scale bar 10 μm.

### Fingers and symmetry breaking

We next sought to identify the source of symmetry breaking in both conditions, with and without application of cyclic uniaxial stretching. It has previously been reported that in MDCK monolayers, cells can migrate tangentially to the monolayer boundary when confined (Doxzen *et al.*, 2013), or perpendicular to the boundary in the form of multicellular groups or fingers during monolayer expansion (Reffay *et al.*, 2011, 2014). During spontaneous elongation of MDCK colonies in the absence of externally applied cyclic stretching (Video 4), we observed that boundary cells tend to move either perpendicularly or tangentially to the colony boundary (Figure 4a). In most of the experiments, an acto-myosin cable, similar to compartmentalization boundaries *in vivo* (Monier *et al.*, 2010; Calzolari, Terriente and Pujades, 2014), was observed in the outer boundary of the colonies at stencil removal (see Figure 4 – figure supplement 1). When this supra-cellular structure is intact, cells at the periphery are reported to undergo clockwise and counter clockwise rotations (Doxzen *et al.*, 2013). In contrast, when a local interruption of this cable appeared, the cell at the vicinity could extend a lamellipodia and move away and radially from the center of the colony (Figure 4 – figure supplement 2a). Apparently, a local defect in the cable could promote the local protrusion of a cell in the direction normal to the edge as shown in laser ablation experiments previously (Reffay *et al.*, 2014). Several local defects could appear within the same colony, thus providing the opportunity for cells in the vicinity to protrude outwards. This cell has been termed leader cell (Reffay *et al.*, 2011) and the collection of cells protruding from the circular colonies along this cell can be identified as the finger-like structures already reported for MDCK monolayers (Reffay *et al.*, 2011, 2014).

**Figure 4.**
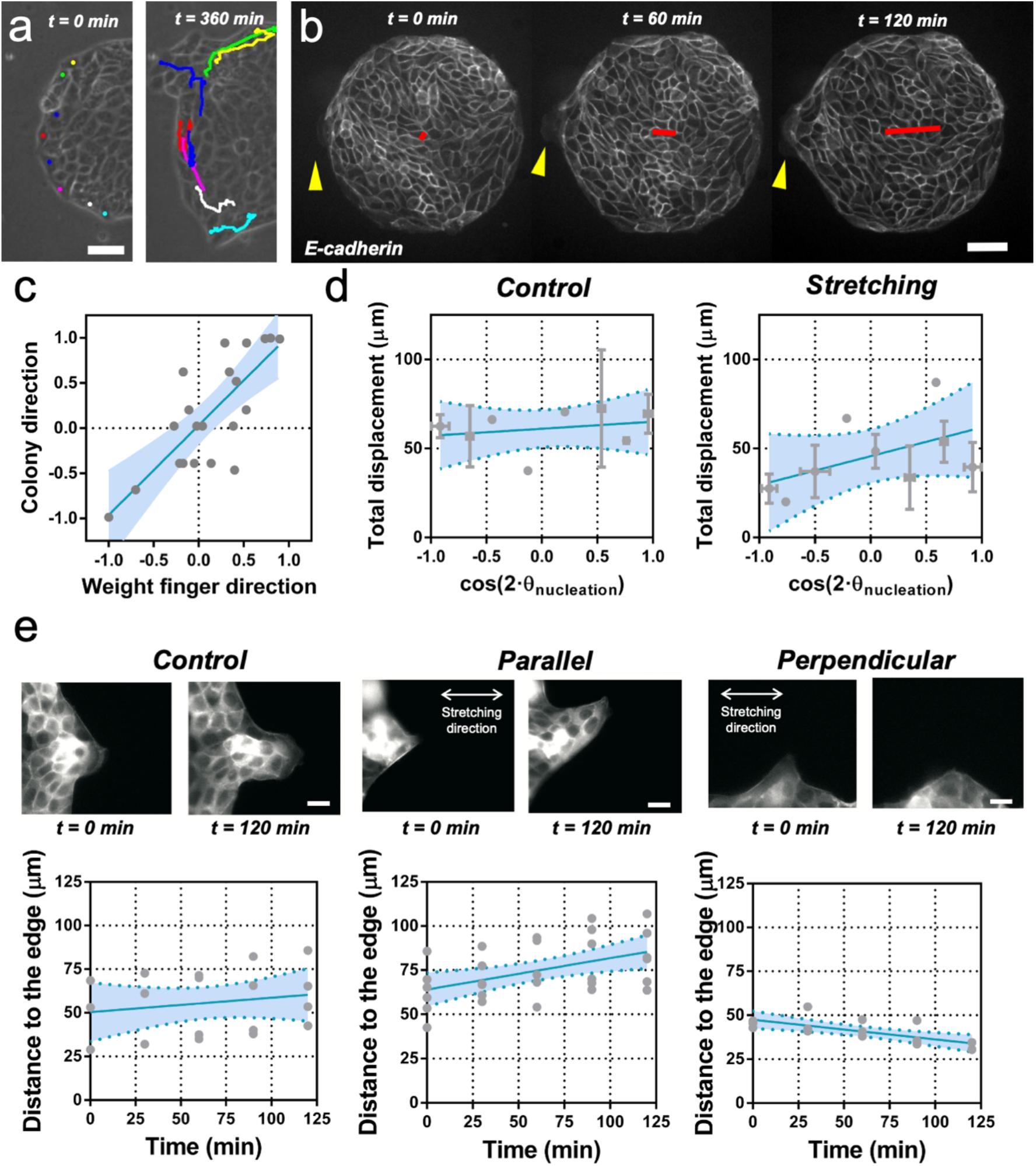
Fingers and symmetry breaking. (**a**) Trajectories of boundary cells during colony expansion. Two types of trajectories are observed: radial and tangential. Scale bar 50 μm. (**b**) Fluorescent images (GFP-E-cadherin) of a colony evolving for 120 min. Red line shows the orientation of the colony according to ellipse fitting (length scales with the change in *Q_xx_*) and yellow arrow indicates a cell migrating radially. Scale bar 50 μm. (**c**) Direction of the colony elongation (quantified by the cosine of 2 times the angle of the main axis of the colony at t = 2 hours) as a function of the weight finger direction (quantified by the average of the cosine of 2 times the angle of each finger trajectory, e.g. the angle corresponding to the vector between the position of the finger at t = 0 hours and t = 2 hours, of each finger weight by the finger’s displacement). N = 5, n_colonies_ = 12 colonies and n_finger_ = 21 fingers. Blue line corresponds to the linear fitting of the data points and the shadowed area corresponds to the 95% confidence interval. Pearson’s correlation coefficient r = 0.7724, p = 0.0001. (**d**) Total displacement of finger growth as a function of its initial position in the colony (angular coordinate from the center of the colony) for control colonies and colonies under uniaxial stretching. N = 5 independent experiments, n_colonies_ = 11 colonies and n_finger_ = 21 fingers (control) and N = 6, n_colonies_ = 10 colonies and n_finger_ = 28 fingers (stretching). Averaged fingers in grey (both position and distance, Mean ± SD), blue line corresponds to the linear fitting of the data points and the shadowed area corresponds to the 95% confidence interval. (**e**) Distance between the tip of the finger and the edge of the monolayer along time, for monolayers in control conditions, stretched parallel and perpendicular to the finger growth direction. N_control_ = 3, N_parallel_ = 4 and N_perpendicular_ = 3 independent experiments and n = 4, 6 and 3 fingers respectively. Individual fingers in grey, blue line corresponds to the linear fitting of the data points and the shadowed area corresponds to the 95% confidence interval.

We performed cell tracking and observed that, on average, these protruding cells are faster than other boundary cells (Figure 4a and Figure 4 – figure supplement 2b and 2c). They are characterized also by radial and directional migrations, in contrast to tangential motion observed in the other cells of the outer region of the colony (Figure 4 – figure supplement 2d and 2e). In general, the motion of these so-called leader cells was directionally persistent and on average the shape of the whole colony followed the same overall direction (Figure 4b and 4c). To correlate colony elongation with leader cell orientation, we analyzed the evolution of a larger number of colonies for 2 hours after stencil removal (Video 4). We quantified the breaking of symmetry by fitting an ellipse to the shape of each colony. We then tracked the positions where finger-like structures were appearing, as well as the direction and distance performed by each of them. We could observe that the elongation direction of the whole colony correlated on average with the direction of the leader cell migration and associated finger (Figure 4c).

We then measured the position and displacement for each finger in control colonies and colonies under cyclic uniaxial stretching (Figure 4d). We observed that, when growing perpendicular to the direction of force application, finger cells performed shorter displacements than when growing parallel to it. In the absence of externally applied cyclic uniaxial stretching, fingers grew a similar amount as when growing parallel the direction of applied uniaxial cyclic stretching and no bias was observed *vis-à-vis* the nucleation position (Figure 4d and Figure 4 – figure supplement 2f). To further explore this effect, we grew MDCK monolayers with straight boundaries either parallel or perpendicular to the external force. Then, we let fingers appear and grow for 2 hours before applying cyclic uniaxial stretching (Figure 4e). When fingers were growing perpendicular to the stretching direction, they shrank upon application of cyclic uniaxial stretching; in contrast, fingers further elongated when parallel to the direction of uniaxial cyclic stretching. Altogether, this suggests that direction of finger-like structures correlates with elongation direction, and that external stretching affects the dynamics of finger growth.

### Collective effects and symmetry breaking

Finger growth correlates with colony elongation. However whether it is a cause or consequence of the symmetry breaking of the shape of the colony remains elusive. We therefore explored the possibility of inducing the growth of fingers and therefore set the direction of elongation of the colonies. Breakage of the acto-myosin cable by laser ablation induces the appearance of leader cells (Reffay *et al.*, 2014). Hence we attempted to trigger the growth of fingers by locally injecting cytochalasin D using a micropipette. The transient injection of this actin polymerization inhibitor was followed by the disruption of the acto-myosin cable (Video 5 and Figure 5 – figure supplement 1). However, the cable reformed, and fingers did not appear. This result shows that breakage of the cable alone doesn’t trigger the growth of fingers in our colonies, and suggests that other mechanisms may be involved.

**Figure 5.**
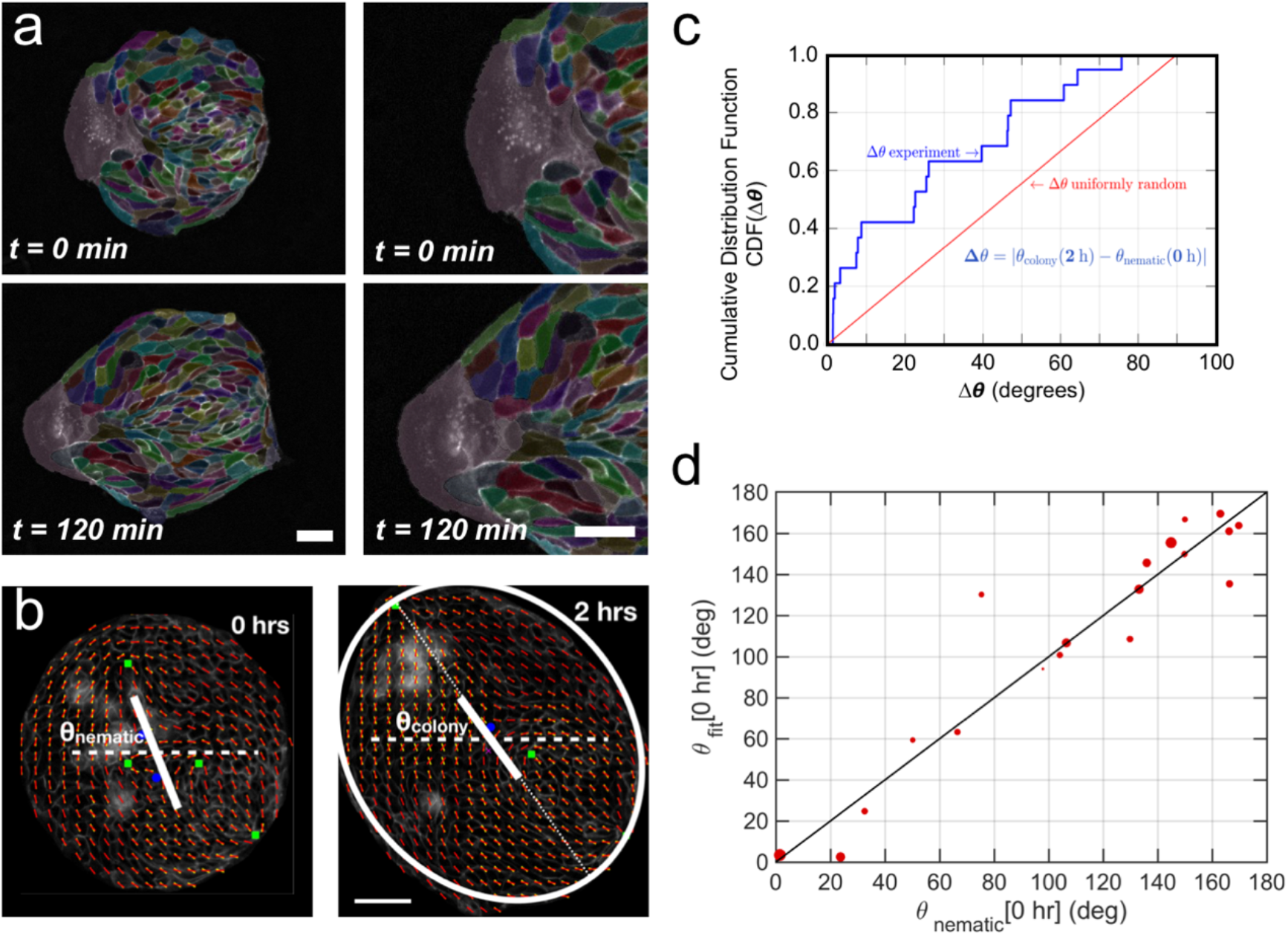
Collective effects and symmetry breaking. (**a**) A leader cell at the boundary of the colony pulls the colony outwards while inner cells deform and elongate. Scale bar 50 μm. (**b**) Cell shape is quantified using a nematic field (red segments). First, the mean cell shape nematic is quantified at the moment of stencil removal (0 hours) and the orientation θ_nematic_ of its mean for the entire colony is obtained. Then, the overall shape of the colony after 2 hours is obtained by fitting an ellipse, whose major axis makes an angle θ_colony_. The yellow directors correspond to fits for the cell shape nematic field obtained with respect to the +½ and −½ topological defects of the experimentally obtained (red) nematic field (also see Supplementary Material Section V for details.) Scale bar 50 μm. (**c**) The cumulative distribution function (CDF) for the difference *Δθ* between θ_nematic_ (0 hours) and θ_colony_ (2 hours) is obtained from n = 19 colonies of N = 5 independent experiments. Red line corresponds to the CDF of a random distribution of the difference *Δθ.* This plot shows a strong correlation between the cell shape nematic and the overall shape symmetry breaking (also see Figure 5 – figure supplement 1d, and Supplementary Material Section IE). (**d**) The experimentally measured angle of mean nematic orientation θ_nematic_ obtained for 19 colonies at t = 0 h is compared with its counterpart θ_fit_ obtained by fitting the experimental data with Eq. 1 of the main paper with respect to the orientation parameter *α* (see Section ID of Supplementary Material). The size of the red circles in (b) is proportional to the magnitude of anisotropy of the colony shape after 2 h. n = 19 colonies of N = 5 independent experiments.

We observed that when a finger moves outward from the colony, cells in the immediate vicinity elongate and seem to reorient their elongation axis towards the finger (Figure 5a). Recent studies have shown that the nematic field of cell elongation and its topological defects could be involved in the growth of bacterial colonies (Doostmohammadi, Thampi and Yeomans, 2016) and in controlling dynamics, death and extrusion of epithelial cells (Kawaguchi, Kageyama and Sano, 2017; Saw *et al.*, 2017; Mueller, Yeomans and Doostmohammadi, 2019). We wondered if the spontaneous elongation of colonies would also be related to the average cell elongation. We followed the evolution of the cell elongation nematic field in different MDCK and MCF 10A colonies during expansion. We first obtained the spatio-temporal cell elongation nematic orientation field *ϕ(x, y, t)* (see Methods) on the experimental time-lapse images (see Figure 5b, Figure 5 – figure supplement 2a-c, Video 6 and Supplementary Material Section IC). We could then obtain the angle *θ*_nematic_ of the average cell-shape nematic field at *t*_0_ which we compared with final colony orientation *θ*_colony_ obtained using the ellipse fitting analysis (Figure 5c, Figure 5 – figure supplement 2d and Supplementary Material Section IC). Strikingly, we observed a clear average cell elongation even at the time of stencil removal *t*_0_, and the corresponding angle *θ*_nematic_ correlated with colony orientation when elongation direction was established for both MDCK and MCF 10A cell lines (Figure 5d, Figure 5 – figure supplement 2d and 2e). The cell elongation orientation field ϕ*(x, y, t)* was not homogeneous at *t*_0_ but exhibited a complex pattern with ±½ topological defects (Figure 5b and Supplementary Material Section ID). Interestingly, an expression that provides equilibrium orientation of liquid crystals with defects and having one constant Frank free energy,

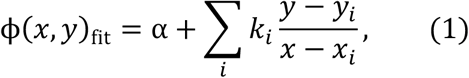

mimics the experimentally observed orientation field *ϕ(x, y)* very well with just one fitting parameter 2 (see Supplementary Material Section ID for details.) Here *k_i_* = ±½ and (*x_i_*, *y_i_*) are the strength and location, respectively, of the topological defects obtained from the experimental image. Thus, the defect position and strength can be used to provide an approximate readout for the orientation of the cell-shape nematic field (Figure 5d). Moreover, the location of finger nucleation seemed to be biased towards the position of topological defects. However, some defect locations were not stable in time and in some cases, the nematic field of cell shapes could only be interpreted in terms of virtual defects outside the colonies, thus suggesting that the mean nematic direction is a better readout for the cell-shape nematic field.

On the one hand, finger nucleation seemed to be correlated with colony elongation direction (Figure 4c). On the other hand, the orientation of tissue elongation correlates with orientation of average cell elongation at *t*_0_ (Figure 5 and Figure 5 – figure supplement 2). This suggests that leader cells moving outward from the colony may not be the cause of symmetry breaking in colony shape, but rather follow from the initial cell shape elongation before stencil removal. Moreover, we found no correlation between breaks of the acto-myosin cable surrounding the colony and the mean nematic direction (Figure 5-figure supplement 4), which suggests that breaks are uniformly distributed along the colony border. We have also shown that breakage of acto-myosin cable after stencil removal, which is associated with leader cell formation (Reffay *et al.*, 2014), did not necessarily induce the growth of fingers in our colonies. Altogether, our results could indicate that the orientation of the mean cell-shape nematic of the colony before stencil removal sets the direction of elongation by triggering the growth of fingers, which appear at the discontinuities of the outer acto-myosin cable located along the nematic orientation, while preventing finger growth at discontinuities located in other directions.

Finally, when looking at the evolution of the mean cell elongation nematic field of colonies under uniaxial cyclic stretching, we observed that it did not change over time (Figure 5 – figure supplement 5). The initial mean direction of cell elongation, either parallel or perpendicular to the external stretching, was maintained throughout 2 hours of external stretching. This suggests that average cell elongation alone does not determine colony elongation direction when subjected to uniaxial cyclic stretching.

### Contributions to symmetry breaking

We next sought to evaluate quantitatively the contribution of cellular processes to elongation. We quantified the contributions of each cellular event using image segmentation, cell tracking and neighbor triangulation (Etournay *et al.*, 2015, 2016) (see Supplementary Material Section IA, Figure 6a and 6b, Figure 6 – figure supplement 1 and Figure 6 – figure supplement 2, and Video 7). This procedure decomposes overall tissue elongation, which is quantified in terms of cumulative total pure shear, into contributions from cell elongation and topological changes. Five main events contribute to total shear: cell elongation change, cell division, cell extrusion, T1 transition, and correlation effects (Etournay *et al.*, 2015, 2016). At the colony scale, shear decomposition plots (Figure 6c and Figure 6 – figure supplement 3) revealed that the total pure shear gives values consistent with elongation estimates from ellipse fitting (Figure 6d). Note that various contributions to shear decomposition exhibit significant variability between experiments (Figure 6 – figure supplement 3). However, we found that after the first 2 hours, the contribution of cell elongation is generally comparable to the total pure shear, with a smaller contribution from other sources (Figure 6d). When looking at the shear decompositions of colonies under cyclic uniaxial stretching (Figure 6e and Figure 6 – figure supplement 3), the cumulative shear values were also similar to the ones obtained by ellipse fitting (Figure 6f). Interestingly, we found however that in that case, shear created by T1 transitions is the main contributor for the total pure shear, while cell elongation does not contribute to tissue elongation (Figure 6f and Figure 6 – figure supplement 3). This indicates that applying oscillatory forces to the tissue changes fundamentally the main mode of tissue elongation by favoring topological rearrangements of the cell network.

**Figure 6.**
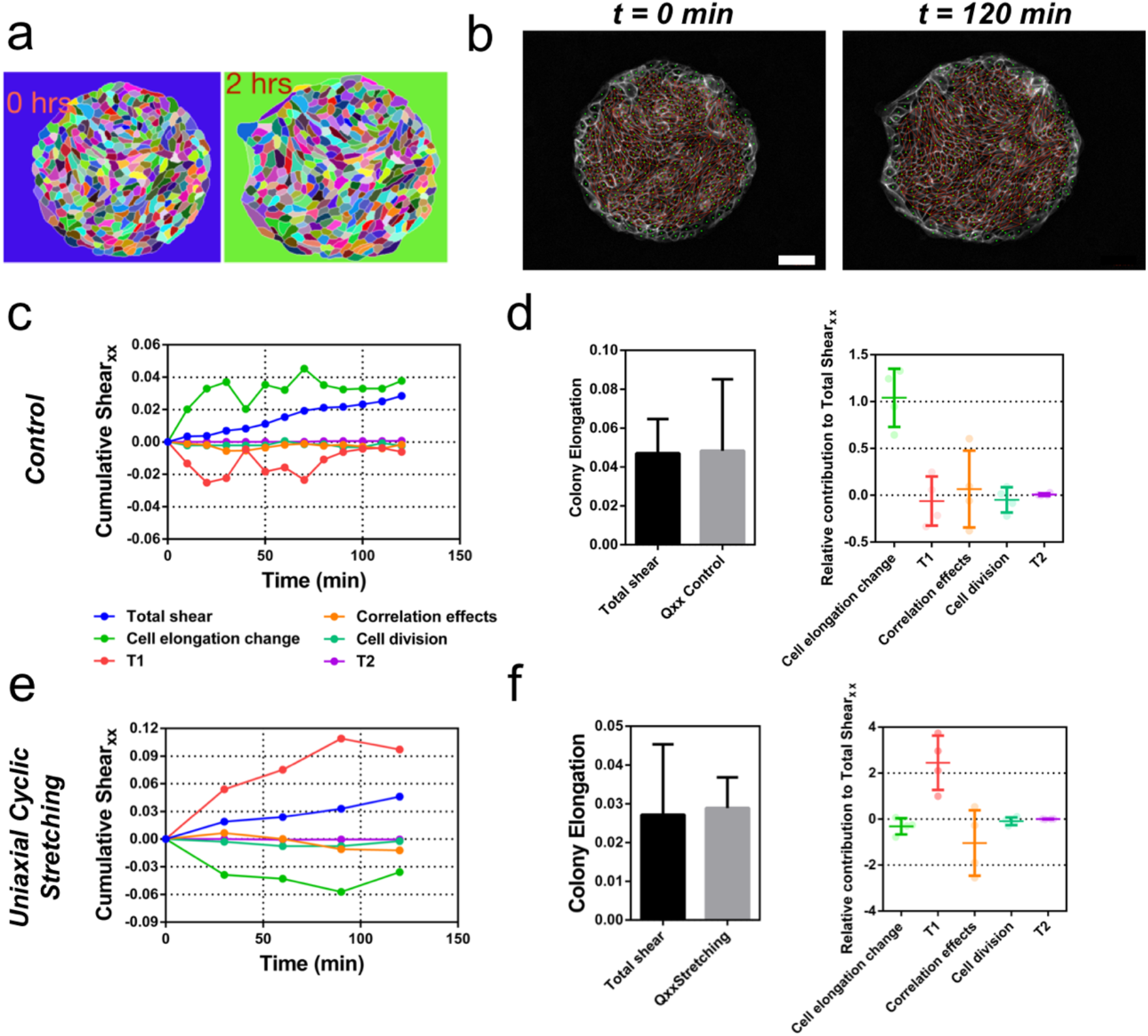
Contributions to symmetry breaking. (**a**) Snapshots of two colonies from control condition segmented and tracked for cells using Tissue-Analyzer (TA) for a duration of *t* = 120 minutes starting from removal of stencil at *t = 0*. Scale bar 50 μm. (**b**) The segmented and tracked images are triangulated in Tissue-Miner (TM). Scale bar 50 μm. The green dots represent the centers of the segmented cells. (**c**) The dynamics of triangulation is analyzed in TM to provide the overall *xx* component of cumulative pure shear strain in a sample colony as a function of time (total shear). (**d**) Comparison between the mean total pure shear obtained from TM and the overall colony pure shear obtained from ellipse fitting (left). Total shear corresponds to n_colonies_ = 4 colonies from N = 2 independent experiments and *Q_xx_* control was obtained from n_colonies_ = 25 colonies of N = 11 independent experiments. Relative contribution of the different processes to the total pure shear (right). Total shear and contributions were obtained from n_colonies_ = 4 colonies from N = 2 independent experiments. (**e**) Cumulative pure shear decomposition for stretched colony. (**f**) Comparison between the mean total pure shear obtained from TM and the overall colony pure shear obtained from ellipse fitting (left) and relative contribution of the different processes to the total pure shear (stretching case). Total shear corresponds to n_colonies_ = 4 colonies from N = 4 independent experiments and *Q_xx_* stretching was obtained from n_colonies_ = 20 colonies of N = 9 independent experiments. Relative contribution of the different processes to the total pure shear (right). Total shear and contributions were obtained from n_colonies_ = 4 colonies from N = 4 independent experiments.

### Vertex model recapitulates symmetry breaking and shear decomposition

We then asked whether a model could reproduce experimental observations of shear decomposition and, in particular, in which conditions tissue elongation would arise from cell elongation or from topological transitions. We developed a vertex model which takes into account mechanical anisotropies such as active stresses and polarized cell bond tension (see Figure 7a and Supplementary Material Section II). We generated a confluent colony of circularly confined cells, in which a unit director **p** that represented the cell polarity was assigned to each cell. Based on orientation of the director, each cell generated an extensile active stress *****σ*****_a_ and bias λ in the base value of its edge contractility to promote cell elongation and active T1 transitions. We assumed that the experimentally measured cell elongation nematic **q** is a readout of the underlying cell polarity **p** (Figure 7b). Hence, the initial spatial distribution of **p** was based on the commonly observed pattern of **q** (Figure 7c). To evolve **p** with time, we imposed that **p** of the exterior cells tended to be parallel to the boundary, whereas the inner cells tended to align their **p** with those of their neighbors (Supplementary Material Section IID).

**Figure 7.**
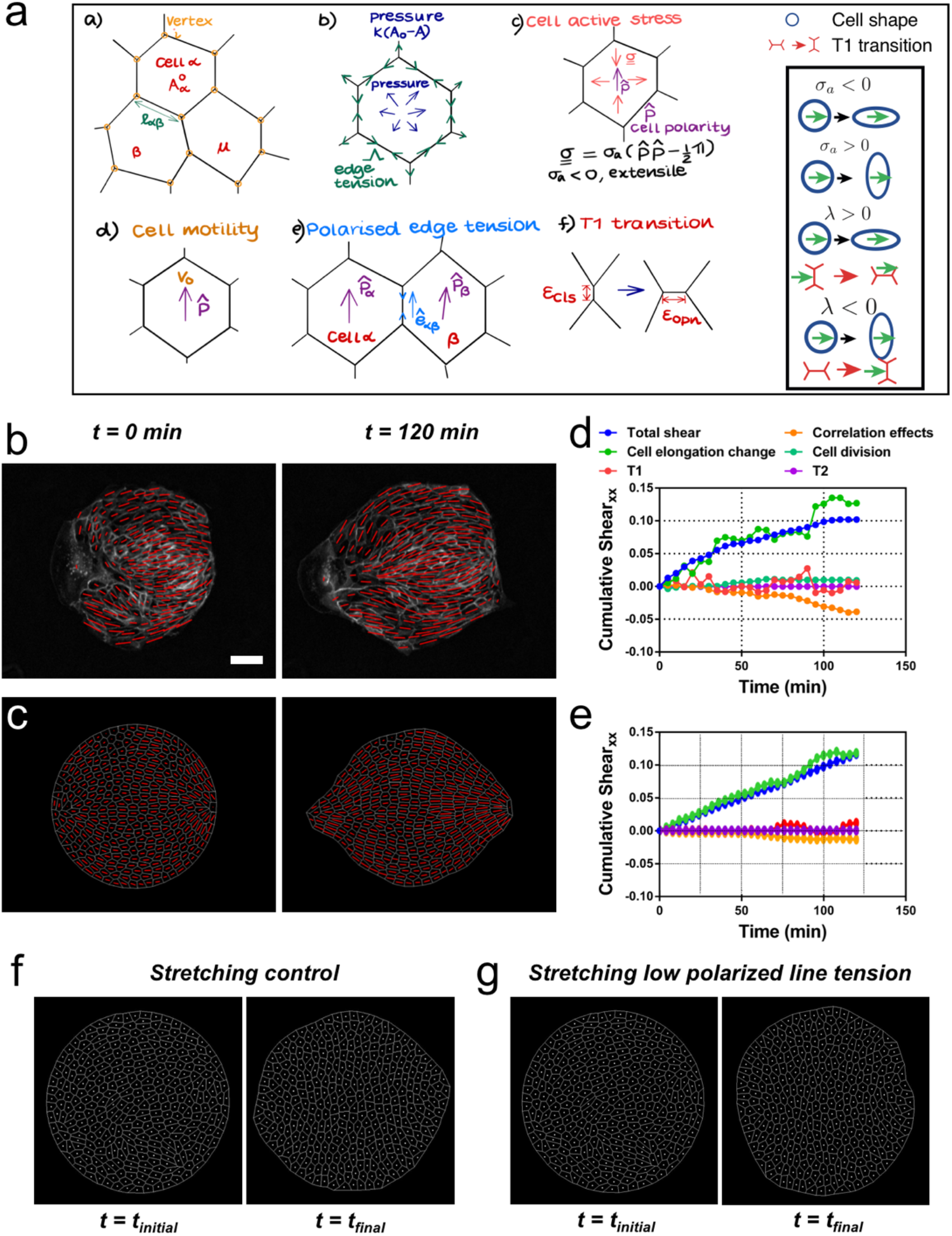
Vertex model recapitulates symmetry breaking and shear decomposition. (**a**) Schematic of vertex model depicting the arrangement of cells, forcing, and topological transitions in a tissue. (a) 2-D monolayer of epithelial cells is represented by polygons, generally sharing a common edge and two vertices between cells. For any cell α shown in the figure, *A*α is the area, 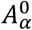 is the preferred area and *l*αβ is the length of the edge shared between cells α and β. The forces on any vertex *i* in the basic vertex model are from (b) pressure due to deviation in the cell area from its preferred value and the tensile force arising from edge or cortical contractility *Λ*. In our model each cell also has a polarity 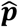 associated with it through which active forces can act on the cell vertices due to (c) anisotropic cell active stress (extensile in our case), (d) cell motility *v_0_* and (e) polarised or biased edge tension that depends on the orientation of the edge 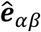 with respect to the polarities of the adjoining cells. (f) When the edge connecting two cells becomes smaller than a critical value ϵ_cls_, the cells are made to modify their neighbours by forming a new edge of length ϵ_opn_ > ϵ_cls_ as shown. Scheme depicting the different possibilities from the model parameters. (**b**) Experimental coarse-grained cell shape nematic at *t=0* and *t=120* min. Scale bar 50 μm. (**c**) A vertex model with internal activity arising from extensile active cell stress (σ_a_′ = −5) and biased edge tension for the cell-cell junctions (λ′ = 20). Prime symbol ′ refers to non-dimensional values (see Supplementary Material Section II). (**d**) Overall *xx* component of cumulative pure shear strain in the sample colony shown in (b) as a function of time (total shear). (**e**) Shear decomposition of the *in silico* experiment which is similar to its experimental counterpart in (**d**). (**f**) A vertex model with an additional aligning term that defines the direction of the uniaxial stretching through cell polarity. When cell-cell junctions are intact, biased edge tension (λ′ = 50) dominates over active cell stress (σ_a_′ = 2) and the colony elongates along the (horizontal) direction of stretch collectively through T1 transitions. (**g**) When the effect of edge tensions is lowered (λ′ = 25) and active cell stress is increased σ_a_′ = 4), the colony elongates perpendicularly to the direction of stretch.

Upon removal of confinement, we found that the simulated colony spontaneously elongates, in a direction set by the orientation of the mean cell elongation nematic field, along with the formation of finger-like structures near the +½ defect, as observed experimentally (see Video 8). Our simulations therefore reproduce experimental observations indicating that colony deformation can be understood without forces exerted by a leader cell at the colony boundary (Figure 7c, Video 8, and Supplementary Material Section II). To test whether we could also recapitulate different contributions of the total pure shear inside the tissue, we performed a shear decomposition analysis of *in silico* movies. We found a qualitatively similar cumulative shear-strain decomposition as was observed in experiments (Figure 7d and 7e), where the main contribution came from cell elongation. Moreover, by changing the relative contributions of the cellular active stress magnitude (*****σ*****_a_) and the edge tension bias (λ), we could modulate the relative contributions from cell elongations and T1 transitions to the total pure shear (Figure 7 – figure supplement 1) as was also observed in experiments with colonies in the absence or presence of cyclic stretching (Figure 6 – figure supplement 3). When *****σ*****_a_ was dominant, the colony elongation was primarily due to cell elongation, whereas when λ was the stronger term, T1 transitions were the main cause of colony elongation. These results reveal possible cellular mechanisms that can govern the process of tissue deformation and influence whether cell elongation, or cellular rearrangements, dominate tissue elongation. Our vertex model assumed that the cell elongation was the main readout for cell polarity, and it did not explicitly account for the effect of substrate stretching. To incorporate uniaxial cyclic stretching we developed further the model. Our results show that initial cell shape elongation does not have a preferential direction (Figure 5 – figure supplement 5), but colony elongation under uniaxial cyclic stretching is along *x* axis, the direction of stretching, and mainly achieved through T1 transitions (Figure 2c and Figure 6f). Also, we report that the elongation happens along *y* axis, perpendicular to the direction of stretching, in single cells and cell colonies with lowered Ecadherin levels (Figure 3a–d). These two experimental observations can be implemented in the model. First, by introducing an additional term ****m****_stretch_ that oriented cell polarisation **p** along *x* axis, for any given initial condition, upon confinement removal at *t*_0_ – this term is inactive in the absence of uniaxial stretching (Supplementary Material Section II). Then, by using cell active stress *****σ*****_a_ > 0, we mimicked the tendency of single cells to elongate perpendicular to the orientation of polarity, *i.e.,* perpendicular to the uniaxial stretching, whereas the bias in the edge tension λ > 0 induced T1 transitions along the polarity of the cell, *i.e.,* parallel to the uniaxial stretching. Thus, in the presence of uniaxial stretching, ****m****_stretch_ oriented cell polarities along *x*, while the relative magnitudes of single cell active stress *****σ*****_a_ and edge-contractility bias *λ* dictated the orientation of the colony elongation. When keeping a lower magnitude of *****σ*****_a_ and a higher value of λ, colonies elongated along *x* through T1 transitions (Figure 7f and Video 9), mimicking colony elongation under uniaxial cyclic stretching (Figure 2 and Figure 6f). On the contrary, by increasing *****σ*****_a_ and lowering λ (Ecadherin deficient colonies), colonies elongate perpendicular to *x* (Figure 7g and Video 9), mimicking colonies under uniaxial cyclic stretching treated with E-cadherin antibody (Figure 3c–d). Therefore, we propose that a competition between the strength of active T1 transitions parallel to the external stretching and active cell stress perpendicular to the external stretching dictate overall colony elongation under uniaxial cyclic stretching. When cell-cell junctions are intact, colony elongation is along the direction of stretching and through T1 transitions (Figure 2c and Figure 6f), suggesting that the tendency of single cells to orient along *y* (Figure 3a–b) is partially screened by cell-cell junctions via T1 transitions. When cell-cell junctions are weakened, active cell stress dominates, and colonies elongate perpendicular to the uniaxial stretching (Figure 3c and 3d), which could be thought of to be closer to a collection of single cells.

### Stretching-dependent elongation is mediated by ROCK

We showed that upon stretching, cells reduced their speed and myosin structures appeared (Figure 3e and 3f). These type of cellular responses to external stretching involve the Rho-associated protein kinase (ROCK) (Hart *et al.*, 2020), which is also involved in cell-cell contacts integrity (Nishimura and Takeichi, 2008; Ewald *et al.*, 2012). We treated MDCK colonies with a ROCK inhibitor (Y-27632 50μM) and followed their behavior for 2 hours, both in the absence and the presence of cyclic uniaxial stretching (Figure 8 – figure supplement 1). When cyclic stretching was applied, elongation along the direction of the uniaxial cyclic stretching was absent (*Q_xx_* ≈ *Q_xy_*) (Figure 8 – figure supplement 1a and 1b). However, colonies still elongated anisotropically, similar to colonies in the absence of application of uniaxial cyclic stretching (Figure 8 – figure supplement 1c). According to our model, when edge tension bias is sufficiently large, the dominant mechanism for tissue elongation switches from single cell elongations to T1 transitions (Figure 7 – figure supplement 1). We observed that uniaxial cyclic stretching triggers this type of elongation (Figure 6f). Therefore, if the effect that application of uniaxial cyclic stretching has at the cellular level was reduced, colonies subjected to cyclic uniaxial stretching should preferentially elongate as if cellular active stress becomes dominant, *i.e.* through single cell elongation. Strikingly, shear decomposition analysis shows that the elongation mechanism shifts from T1 transition-dominant to cell elongation-dominant when colonies under uniaxial cyclic stretch have ROCK inhibited (Figure 8a–b). In summary, colonies under cyclic uniaxial stretching elongate through T1 transitions rather than through cell elongation (red dot in Figure 8c) similar to what our model predicts for colonies with increased biased edge tension. In contrast, colonies elongate spontaneously largely through single cell elongation, which the model predicts when the effect of the cellular active stress is more dominant (green dot in Figure 8c). Strikingly, the application of a ROCK inhibitor leads to single cell elongation dominating over T1 (orange dot in Figure 8c), effectively suppressing the effect of uniaxial cyclic stretching on the mode of colony deformation (Figure 8 – figure supplement 1).

**Figure 8.**
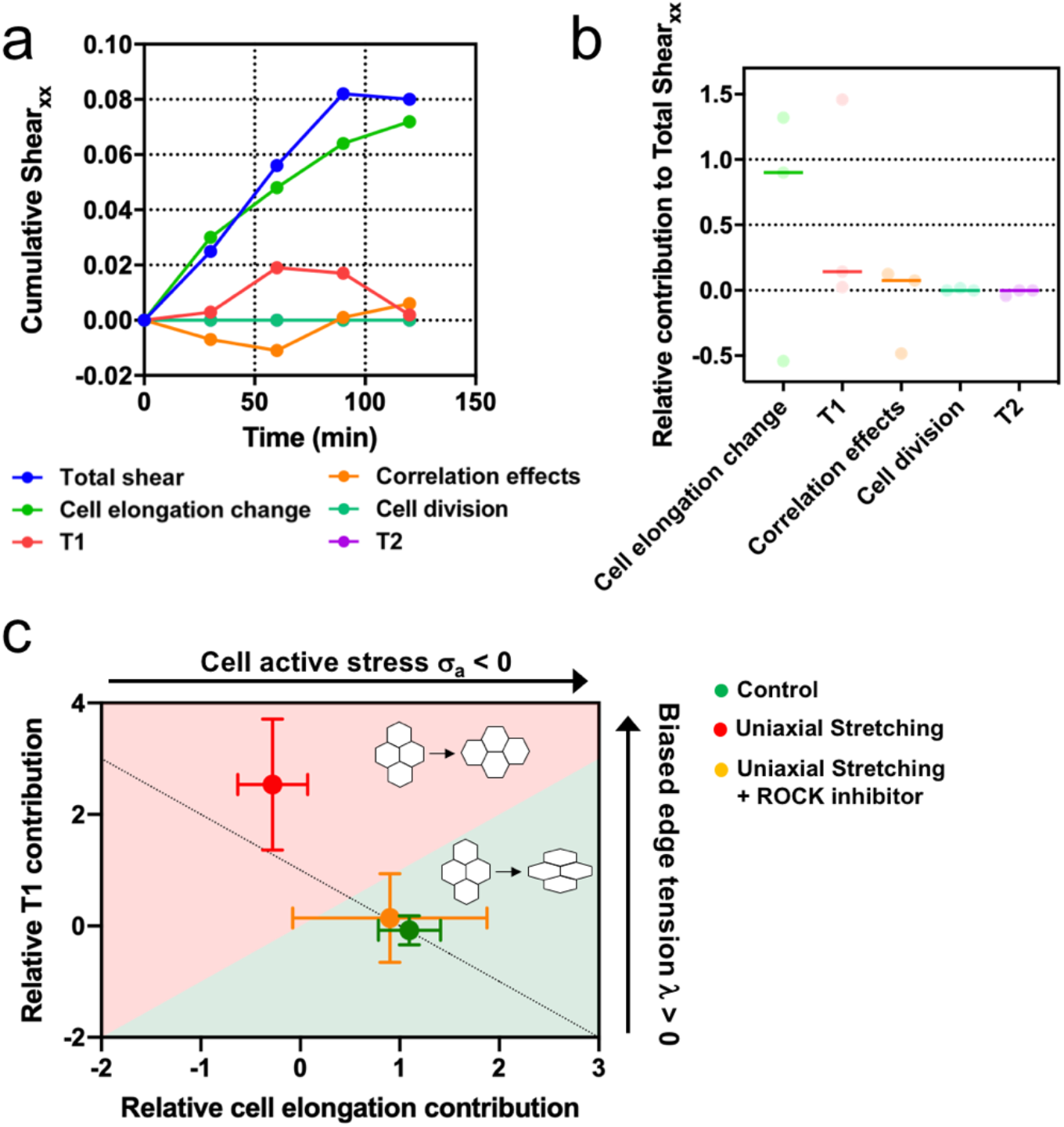
Stretching-dependent elongation is mediated by ROCK. (**a**) Cumulative pure shear decomposition for stretched colony in the presence of ROCK inhibitor Y-27632 50 μM. (**b**) Relative contribution of the different processes to the total pure shear. Total shear and contributions were obtained from n_colonies_ = 3 colonies from N = 3 independent experiments. Individual experiments and median are plotted. (**c**) Contribution of single cell elongation to total elongation versus contribution of T1 transitions to total elongation. In the green area single cell elongation dominates over T1, whereas red area corresponds to T1 transitions dominating over cell elongation. Along the dashed line, contribution of correlation effects and oriented cell division to total shear is close to zero. Above this line this contribution is negative and below the line is positive. According to our vertex model, by changing the strength of cellular active stress and biased edge tension, colonies change the relative contribution of each mechanism of elongation. Experiments show that epithelial colonies spontaneously elongate through single cell elongation (green point, N = 2, n_colonies_ = 4), whereas colonies under uniaxial stretching elongate through T1 (red dot, N = 4, n_colonies_ = 4). When ROCK activity is inhibited, colonies under uniaxial stretching elongate through single cell elongation (orange dot, N = 3, n_colonies_ = 3). Points represent median ± SD.

## Discussion

Tissue spreading is key during embryonic development (Bénazéraf *et al.*, 2010; Campinho *et al.*, 2013; Etournay *et al.*, 2015). Epithelial cells migrate in a collective and cohesive manner. In many cases symmetry is broken leading to shape transformation. The resulting tissue kinematics and the underlying mechanisms for this symmetry breaking has been studied using original approaches, and is understood with a variety of cell-based and continuum models *in vivo* (Blanchard *et al.*, 2009; Clarletta, Ben Amar and Labouesse, 2009; Aigouy *et al.*, 2010; Etournay *et al.*, 2015; Guirao *et al.*, 2015; Alt, Ganguly and Salbreux, 2017). On the *in vitro* situation too, combination of theory and experiments have been applied on similar phenomena (Mark *et al.*, 2010; Tarle *et al.*, 2015; Kawaguchi, Kageyama and Sano, 2017; Saw *et al.*, 2017).

In the present work, we sought to integrate knowledge from *in vitro* and *in vivo* studies to test new ideas for breaking symmetry through collective effects. First we showed that initially circular colonies of three different epithelial cell lines spontaneously expanded in a non-isotropic manner and the elongation observed was similar in magnitude to *Drosophila* wing blade elongation *in vivo* (Etournay *et al.*, 2015). However, the undeformed circular colonies have a non-zero average cell elongation even before spreading starts, which determines orientation of the final colony shape. Our analysis also showed that the cell orientation patterns are not homogeneous but spatially organized and directed by the presence of ±½ topological defects. It was already shown that topological defects in the cell elongation nematic field have a key role on epithelial dynamics (Kawaguchi, Kageyama and Sano, 2017) and on cell death and extrusion (Saw *et al.*, 2017). Our results reinforce the idea that cell elongation nematic field, which could only arise from collective interaction between cells, can have an impact on epithelial morphogenesis.

Since developing tissues are regularly subjected to internal oscillations (Solon *et al.*, 2009; He *et al.*, 2010; Rauzi, Lenne and Lecuit, 2010) and external pulsatory forces (Zhang *et al.*, 2011; Hayes and Solon, 2017) in a number of living organisms, we explored the effect of external forces in the *in vitro* circular colonies. We observed that the direction of elongation could be rectified by imposing an external uniaxial cyclic force. This particular behavior is of great interest from an *in* vivo point of view: cyclic contraction could direct elongation in specific directions to trigger the next steps in morphogenesis. The generic localizations of muscles connected to epithelial layer (Zhang and Labouesse, 2012) could have this essential function for tissues which would otherwise elongate in any direction like in our assay. Finally, direction of this tissue rectification is along the external force, but perpendicular to the reorientation of single cell under external uniaxial stretching, and this further supports the collective nature of the phenomenon.

We also systematically quantified the shear deformation kinematics of the colony and demonstrated that some of the colonies exhibit shear decomposition patterns similar to those observed during *Drosophila* pupal wing elongation (Etournay *et al.*, 2015). Moreover, while the colony deformation for the control colonies was dominated by cell elongation, T1 transitions were the main drivers of the colony shape anisotropy under cyclic uniaxial stretching thus indicating different mechanisms at work. Thus, the current work thus makes a direct comparison and contrast tissue kinematics between *in vivo* and *in vitro* cases.

Finally, we developed a vertex model that takes into account mechanical anisotropies. Cell anisotropic active stress arising in the cell core, cortical contractility at the cell-cell junctions and cell motility are three of the important forces involved in morphogenesis of epithelial monolayers. To our knowledge, this is the first attempt to systematically show how each of these activities acts on tissue kinematics. Specifically, we showed that cell anisotropic active stress results mainly in cell elongation, whereas anisotropies in cortical contractility primarily effects cell intercalations or T1 transitions. Including cell motilities appears to enhance tissue shear generated by the other two modes of internal forcing. We perturbed active stress and line edge tension by inhibiting ROCK, which has been reported to be involved in cell-cell contact integrity *in vivo* (Nishimura and Takeichi, 2008; Ewald *et al.*, 2012), and recently, in cell responses to stretching *in vitro* (Hart *et al.*, 2020). This led to experimentally blocking the ability of colonies to respond to the externally applied uniaxial cyclic stretching. By doing so, colonies which primarily elongate through cell intercalations, shifted to a single cell elongation driven mechanism.

From our simulations, we could demonstrate that symmetry breaking and finger formation in colonies and the corresponding tissue kinematics observed in our experiments could be brought about by collective active behavior of the colony cells and does not require special action of leader cells (Theveneau and Linker, 2017). This result echoes experiments in which the emergence of the leader cells and the fingering behavior at the border were suggested to arise due to the internal stress pattern in the tissue (Vishwakarma *et al.*, 2018). On the other hand, there are many excellent models in which the boundary cells are ascribed special motility properties that could replicate the experimental results on border fingering (Mark *et al.*, 2010; Tarle *et al.*, 2015). Thus, although leader cells at the boundary may play a role in the border fingering, our experimental findings and simulations clearly indicate that the cell-level internal activities and cell-cell interactions are sufficient to cause symmetry breaking in the colony shape and its overall kinematics via the collective cell-shape nematic field.

## Conclusion

Our results show that cell elongation nematic field can have an impact on epithelia morphogenesis. It was already reported that topological defects in the cell elongation nematic field have a key role on epithelial dynamics (Kawaguchi, Kageyama and Sano, 2017) and on cell death and extrusion (Saw *et al.*, 2017). Now, we showed that circular epithelial colonies when in confinement build up a mean nematic orientation. This symmetry breaking results from the inner activity of cells, and sets the direction for colony elongation. Epithelia changes in shape could be revisited *in vivo* with this new framework, leading to potential generic rule of morphogenesis in developmental biology.

## Materials and Methods

### Cell culture

MDCK cells (GFP-E-cadherin strain (Adams *et al.*, 1998), GFP-Actin strain, mCherry-actin / GFP-myosin strain (Klingner *et al.*, 2014)) were cultured in Dulbecco’s modified Eagle medium (D-MEM) 1g/l glucose (Invitrogen), supplemented with 10% fetal bovine serum (FBS) (Invitrogen) and 1% penicillin-streptomycin (and the corresponding resistance for each strain). Cells were passaged every 2-3 days using standard procedures. Caco-2 cells (ATCC) were cultured in minimum essential media (MEM) supplemented with Earle’s salts (Life Technologies), 20% fetal calf serum (FCS) (Invitrogen), 0.1 mM non-essential amino acids, 1 mM sodium pyruvate and 40 μg/ml Gentamicin at 37°C and 5% CO_2_. Culture was passaged every 3-4 days. MCF 10A cells (ATCC) were cultured in Dulbecco’s modified Eagle medium (D-MEM) 1g/l glucose (Invitrogen), supplemented with 10% horse serum (Invitrogen), 5 μg/ml insulin, 40 μg/ml Gentamicin, 2 mM L-Glutamine, 0.5 μg/ml Hydrocortisone and 2 ng/ml human epidermal growth factor (hEGF). Cells were passaged every 2-3 days. Cells tested negative for mycoplasma.

### Fabrication of PDMS membranes and stencils

Poly(dimethylsiloxane) (PDMS) (Sylgard) was prepared by mixing the pre-polymer and the cross-linker at a 10:1 ratio. To prepare stretchable membranes, uncured PDMS was first centrifuged (3000 rpm for 5 min) to remove air bubbles. Afterwards, the PDMS was spin-coated on a flat polystyrene (PS) surface (500 rpm for 1 min) and cured at 65°C overnight. PDMS stencils were prepared as described previously (Ostuni *et al.*, 2000). Briefly, SU-8 2050 molds containing circles of 250 μm in diameter were prepared by standard photolithography. Uncured PDMS was then spin-coated on molds to a thickness lower than the height of the microstructures (50 μm) and cured overnight at 65°C. Stencils for the finger experiments were prepared by spin-coating uncured PDMS on a flat surface.

### Cell seeding on stencils

The PDMS stencils were cut, peeled off the mold, and placed in a 2% Pluronic F-127 (Sigma-Aldrich) in PBS for 1 h. The stencils were then kept in PBS for 2h and dried under the hood flow. PDMS stretchable membranes were cut and then activated using O_2_ plasma. The membranes and the stencils were exposed to UV light in the cell culture hood for 10 min. Afterwards, stretchable membranes were incubated with fibronectin 20 μg/ml for 1 h, rinsed with PBS and dried. PDMS membranes were placed on a PS holder, and the PDMS stencils were deposited on top of the membrane right after. A rectangular PDMS chamber was attached onto the membrane using vacuum grease, and cells were seeded at a density of 20000 cells/mm^2^ (Serra-Picamal *et al.*, 2012) for 4 h. When cells were attached, the medium was changed and the membrane with the cells was kept in the incubator. Local cell density could vary within each colony. We followed the dynamics of assembly of the colony prior removal of the stencil and we could see that cellular clusters size distribution and respective location within the cavity at plating could contribute to these variations. Once they formed confluent circular colonies, the stencils were removed with tweezers carefully before starting the experiment. Some of the colonies exhibited elongation in the short time window between stencil removal and the start of image acquisition.

### Time-lapse microscopy

After stencil removal, the medium was replaced by L-15 (Leibovitz’s L-15 medium, Invitrogen) supplemented with 10% FBS. Cells were then observed under a Nikon Ti inverted microscope using either a x10 or a x20 objective for 6 h at 37°C. Images were acquired every 5 min.

### Stretching experiments

The stretching device was designed in the lab. Briefly, a Thorlabs motor (Thorlabs z812 motor, Thorlabs) was controlling the motion of a PDMS membrane, and everything was mounted on a custom-made microscope stage. Circular colonies were plated on PDMS membranes, and after removal of the stencils, samples were placed on the microscope. Cyclic uniaxial stretches were applied and images were taken every 30 minutes typically shortly to prevent interfering with the time course of the experiments. We probed 3 times for cycles, 20s, 60s, 120s, and 3 extensions, 5%, 10%, and 20%. The shape of the cycles was triangular. We checked that the PDMS stencils were elastic at all extension and frequency ranges.

### Chemical treatments

To prevent the formation of E-cadherin-mediated adhesion, cells were incubated for 30min with L-15 medium containing 5 mM EDTA (Sigma-Aldrich) and 10 μg/ml anti-E-cadherin blocking antibody that targeted the extracellular domain of E-cadherin (Gumbiner, Stevenson and Grimaldi, 1988) (uvomorulin, monoclonal anti-E-cadherin antibody, Sigma); after incubation, the medium was replaced by normal L-15 and the experiment started. The inhibition of ROCK was done by incubating cells with Y-27632 50 μM solution (Sigma-Aldrich) from 30 minutes before the experiment started until the end of the experiment.

### Finger dynamics experiments

For the finger test after growth, we let finger grow for 2 hours, and we subsequently applied the cyclic stretch.

### Colony shape change analysis

Shape change analysis was performed using ImageJ (http://rsb.info.nih.gov/ij, NIH). The outline of the colony on phase contrast images was ellipse fitted at every time point. Major axis *a*, minor axis *b*, and ellipse orientation*θ* were obtained. We computed ****Q****, defined as *Q_xx_(t)* = ½ ln(*a(t)/b(t)*)·cos*2·*(*θ(t)* − *α*) and *Q_xy_(t) =* ½ln(*a(t)/b(t)*)·sin*2·*(*θ(t)* − *α*) being *α* = *θ*(t_final_) to quantify cell colony elongation. In uniaxial stretching experiments, the *x* axis corresponds to the direction of the external stretch and ****Q**** components are defined as *Q_xx_(t) =* ½ ln(*a(t)/b(t)*)·cos*2·*(*θ(t)* − α) and *Q_xy_(t) =* ½ln(*a(t)/b(t)*)·sin*2·*(*θ(t)* − α) being *α* = *0*.

### Velocity analysis

The centroid trajectories of cells were tracked using the manual tracking plug-in in ImageJ. Data analysis was performed using a custom-made code in MATLAB (The MathWorks). Cell positions were characterized by a vector **r** (t), with t denoting time and **r** position in space (bold letter refers to a vector). Every recorded cell position during the time-lapse experiment was defined as **r**_i_ = **r**(t_i_), where t_i_ = iΔt are the times of recording and Δt denotes the duration of time-lapses. The average velocity of each cell was then defined as v = (1/T)·∑_i_r_i_, being r_i_ the module of the vector **r**_i_ and T the total duration of the trajectory.

### Colony segmentation and cell tracking

Movies acquired using a MDCK GFP-E-cadherin strain were first pre-processed with FIJI. The *subtract background* function was applied to remove noise. Images were then loaded to Tissue Analyzer (TA) v8.5 (Aigouy *et al.*, 2010) for edge detection and cell tracking.

### Orientation field of the cells and topological defects

First, the background noise of the time-lapse images of the elongating MDCK colonies was reduced with the *subtract background* function by using a rolling ball radius of 40 px. The resulting images were then subjected to *band-pass* filter with upper and lower limits of 40 px and 3 px, respectively. The background noise from this output was reduced by using the *subtract background* command again with a rolling ball radius of 40 px. The processed images from each experiment were analyzed with the OrientationJ plugin of FIJI to quantify their local spatial orientation that reflects the underlying cell elongation. Within this plugin we used a local smoothing window of 20 px (approximately of the size of the cells) to obtain the *structure tensor* at discrete points on a grid of 20 px × 20 px. The plugin provides the dominant direction ϕ_*i*_ of the structure tensor at each point *(x_i_, y_i_)* that represents the local *orientation* field *q_i_* = *cos ϕ_i_ e_1_* + *sin ϕ_1_ e_2_* for cell elongation. The OrientationJ plugin also provides the *coherence C* of the structure tensor to quantify the strength of the orientation; *C* ≈ 0 and *C* ≈ 1 would approximately correspond to rounded and elongated cells, respectively. The orientation or the director field **q** thus obtained was further quantified by studying the spatiotemporal evolution of ±½ topological defects that were obtained by calculating the winding number over unit-cells of the underlying grid. The local smoothing window of 20 px for obtaining the *structure tensor*, which is approximately of the size of cells, ensured that the most robust defects were observed. The validity of this procedure was cross-verified with the smoothed cell-shape nematic field and the corresponding ±½ defects from the segmented and triangulated data of the experiments processed in Tissue Miner (TM) (Figure 5 – figure supplement 1 and Supplementary Material Section I). Finally, the orientation of mean cell-shape nematic calculated at 0 hr was compared with shape orientation of the colony at *t* = 2 hrs. For obtaining the cell-shape nematic field for MCF 10A colonies, the procedure was the same as for MDCK control but the numerical parameters used were, rolling ball radius for *subtract background* 50 px, no band pass filter, and local smoothing window of 15 px and grid size of 30 px for OrientationJ. Similarly, for obtaining the cell-shape nematic field for stretched colonies, the procedure was the same as for MDCK control but the numerical parameters used were, rolling ball radius for *subtract background* 50 px, no band pass filter, and local smoothing window of 40 px and grid size of 30 px for OrientationJ. See Supplementary Material Section I for more details (also see Video 6).

### Acto-myosin cables

In order to image acto-myosin supracellular cables at the boundary of colonies either cells expressing mCherry-actin / GFP-myosin strain or cell immunostaining were used. To disrupt acto-myosin cables, local injection of 4 μM Cytochalasin D using a micropipette was performed. Glass micropipettes were connected to a microinjection system (CellTram vario, Eppendorf). The position of the pipette tip was controlled in *x, y, z* by using a micromanipulator. The system was mounted on an epifluorescence inverted microscope to record the process. Cytochalasin D was released locally for about 10 min. To detect discontinuities in the cable, fluorescent images of actin and myosin were treated using Fiji as follows. First, a median filter was applied and the background subtracted. Then, a contrast-limited adaptive histogram equalization (CLAHE) was used and the background was again removed using an exponential function. Images corresponding to actin and myosin were added. Finally, a Laplacian of Gaussian filter was applied to the resulting image. Once the cable structure was revealed, the positions of the defects were identified.

### Quantification of cellular deformations, topological transitions and their contribution to pure shear deformation

After obtaining the geometrical and topological information of the colonies from the series tracked images generated using TA, TM was used to extract, triangulate and store the data with the help of an automated workflow. The database obtained after this stage of analysis was used to quantify various state properties such as cell area, neighbor number, cell elongation and the contribution of different cellular processes to deformation using scripts written both in *R* and in *Python*. TM was further used to quantify the contributions of various cellular events such as cell elongation and topological transitions to the colony deformation. More details about this analysis can be found in Supplementary Material Section I (also see Video 7).

### Vertex Model Simulations

A vertex model was developed with an addition of a unit nematic director **p** to every cell. The orientation of the boundary cell **p** was maintained parallel to the boundary, whereas **p** for internal cells were modeled to tend to align with the **p** of its neighbors. In these simulations, an extensile active stress 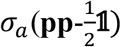 with σ_a_<0 acts to increase cell elongation along **p**. In addition, a bias *λ* was also applied on the basic edge tension with respect to the director **p** of its neighboring cells. For positive *λ*, this bias reduces (increases) the tension of the edge along (perpendicular to) **p**. Consequently, the closure (formation) of edges is enhanced in the direction perpendicular (parallel) to **p**. Hence, the T1 transitions in the region around the cells are oriented to cause shear elongation (contraction) along (perpendicular) to **p**. The colony is provided with an initial condition for **p** that mimics the initial configuration of experimentally frequently observed cell shape nematic fields **q** with two +½ defects that are separated by a distance. The initial polar vector orientation along the nematic axis are chosen at random such that the total polarity is zero, and the dynamics of polarity reorientation is independent on a **p** → −**p** flipping of the polarity axis. To begin with, the cell positions and director orientations were evolved under colony confinement until cell position and **p** do not change significantly. The confinement is then removed to see how the colony breaks symmetry in its shape. In another set of simulations, a small motility *v_0_* was also provided to the internal cells (Figure 7 – figure supplement 1). Similar to the experiments, the output of these simulations was also processed in TM and analyzed for topological defects and pure shear decomposition. For colonies subjected to uniaxial cyclic stretching, **p** for any cell was provided with an additional tendency to align along the stretching orientation (*x-*axis). Moreover, the active stress in this case σ_a_ > 0 was contractile, i.e., the cell tended to elongate perpendicular to the orientation of **p**. More details of the simulations are provided in Supplementary Material Section II (also see Videos 8 and 9).

### Statistics

No statistical methods were used to predetermine sample size. Measurements were performed on different colonies (n) obtained in different independent experiments (N). The exact values are defined at the caption of each figure. Data presentation (as Mean value ± standard deviation (SD) or as Mean value ± standard error of the mean (SE)) is defined at the corresponding figure caption. D’Agostino normality test, Mann-Whitney test and Pearson’s correlation were performed using GraphPad Prism 8. Specific values are noted at the corresponding figure captions.

## Acknowledgments

We acknowledge M. Popovic and R. Etournay for stimulating discussions and help with Tissue Miner, J. Nelson, P. Silberzan, S. Coscoy for sharing the MDCK strains, and J. Prost, K. Dayal and M. Zapotocky for helpful suggestions, D. Rodriguez, M. Lieb and the Riveline Lab. D.R. and M.M.I. thank MPI-PKS and MPI-CBG for hospitality and many interactions. D.R. acknowledges support from CNRS (ATIP), ciFRC Strasbourg, the University of Strasbourg, Labex IGBMC, Foundation Cino del Duca. This study with the reference ANR-10-LABX-0030-INRT has been also supported by a French state fund through the Agence Nationale de la Recherche under the frame programme Investissements d’Avenir labelled ANR-10-IDEX-0002-02. M.M.I. acknowledges funding from Department of Biotechnology, Government of India (BT/06/IYBA/2012) and Industrial Research and Consultancy Center (IRCC), IIT Bombay. G.S. was supported by the Francis Crick Institute which receives its core funding from Cancer Research UK (FC001317), the UK Medical Research Council (FC001317), and the Wellcome Trust (FC001317).

## References

Adams, C. L. et al. (1998) ‘Mechanisms of Epithelial Cell–Cell Adhesion and Cell Compaction Revealed by High-resolution Tracking of E-Cadherin– Green Fluorescent Protein’, The Journal of Cell Biology, 142(4), pp. 1105–1119. doi: 10.1083/jcb.142.4.1105.

Aigouy, B. et al. (2010) ‘Cell Flow Reorients the Axis of Planar Polarity in the Wing Epithelium of Drosophila’, Cell, 142(5), pp. 773–786. doi: 10.1016/j.cell.2010.07.042.

Alt, S., Ganguly, P. and Salbreux, G. (2017) ‘Vertex models: From cell mechanics to tissue morphogenesis’, Philosophical Transactions of the Royal Society B: Biological Sciences, 372(1720), p. 20150520. doi: 10.1098/rstb.2015.0520.

Bénazéraf, B. et al. (2010) ‘A random cell motility gradient downstream of FGF controls elongation of an amniote embryo.’, Nature, 466(7303), pp. 248–252. doi: 10.1038/nature09151.

Blanchard, G. B. et al. (2009) ‘Tissue tectonics: morphogenetic strain rates, cell shape change and intercalation’, Nature Methods. Nature Publishing Group, 6(6), pp. 458–464. doi: 10.1038/nmeth.1327.

Calzolari, S., Terriente, J. and Pujades, C. (2014) ‘Cell segregation in the vertebrate hindbrain relies on actomyosin cables located at the interhombomeric boundaries’, EMBO Journal, 33(7), pp. 686–701. doi: 10.1002/embj.201386003.

Campinho, P. et al. (2013) ‘Tension-oriented cell divisions limit anisotropic tissue tension in epithelial spreading during zebrafish epiboly.’, Nature cell biology. Nature Publishing Group, 15(12), pp. 1405–14. doi: 10.1038/ncb2869.

Clarletta, P., Ben Amar, M. and Labouesse, M. (2009) ‘Continuum model of epithelial morphogenesis during caenorhabditis elegans embryonic elongation’, Philosophical Transactions of the Royal Society A: Mathematical, Physical and Engineering Sciences, 367(1902), pp. 3379–3400. doi: 10.1098/rsta.2009.0088.

Doostmohammadi, A., Thampi, S. P. and Yeomans, J. M. (2016) ‘Defect-Mediated Morphologies in Growing Cell Colonies’, Physical Review Letters, 117(4), pp. 1–5. doi: 10.1103/PhysRevLett.117.048102.

Doxzen, K. et al. (2013) ‘Guidance of collective cell migration by substrate geometry’, Integrative Biology, 5, pp. 1026–1035. doi: 10.1039/C3IB40054A.

Etournay, R. et al. (2015) ‘Interplay of cell dynamics and epithelial tension during morphogenesis of the Drosophila pupal wing’, eLife, 4(JUNE2015), pp. 1–51. doi: 10.7554/eLife.07090.

Etournay, R. et al. (2016) ‘TissueMiner: A multiscale analysis toolkit to quantify how cellular processes create tissue dynamics’, eLife, 5(MAY2016), pp. 1–28. doi: 10.7554/eLife.14334.

Ewald, A. J. et al. (2012) ‘Mammary collective cell migration involves transient loss of epithelial features and individual cell migration within the epithelium.’, Journal of cell science, pp. 2638–2654. doi: 10.1242/jcs.096875.

Faust, U. et al. (2011) ‘Cyclic stress at mHz frequencies aligns fibroblasts in direction of zero strain.’, PloS one, 6(12), p. e28963. doi: 10.1371/journal.pone.0028963.

Fodor, É. et al. (2018) ‘Spatial Fluctuations at Vertices of Epithelial Layers: Quantification of Regulation by Rho Pathway’, Biophysical Journal, 114(4), pp. 939–946. doi: 10.1016/j.bpj.2017.12.026.

Gilmour, D., Rembold, M. and Leptin, M. (2017) ‘From morphogen to morphogenesis and back’, Nature, 541(7637), pp. 311–320. doi: 10.1038/nature21348.

Guillot, C. and Lecuit, T. (2013) ‘Mechanics of Epithelial Tissue Homeostasis and Morphogenesis’, Science, 340(6137), pp. 1185–1189. doi: 10.1126/science.1235249.

Guirao, B. et al. (2015) ‘Unified quantitative characterization of epithelial tissue development’, eLife, 4(DECEMBER2015), pp. 1–52. doi: 10.7554/eLife.08519.

Gumbiner, B., Stevenson, B. and Grimaldi, a. (1988) ‘The role of the cell adhesion molecule uvomorulin in the formation and maintenance of the epithelial junctional complex.’, Journal of Cell Biology, 107(4), pp. 1575–1587. doi: 10.1083/jcb.107.4.1575.

Harris, A. R., Daeden, A. and Charras, G. T. (2014) ‘Formation of adherens junctions leads to the emergence of a tissue-level tension in epithelial monolayers.’, Journal of cell science, pp. 2507–2517. doi: 10.1242/jcs.142349.

Hart, K. C. et al. (2020) An Easy-to-Fabricate Cell Stretching System Reveals Density-Dependent Mechanical Regulation of Collective Cell Movements in Epithelial Homeostasis. doi: https://doi.org/10.1101/2020.08.24.265629.

Hayes, P. and Solon, J. (2017) ‘Drosophila dorsal closure: An orchestra of forces to zip shut the embryo’, Mechanisms of Development. The Authors, 144, pp. 2–10. doi: 10.1016/j.mod.2016.12.005.

He, L. et al. (2010) ‘Tissue elongation requires oscillating contractions of a basal actomyosin network.’, Nature cell biology, 12(12), pp. 1133–42. doi: 10.1038/ncb2124.

Kawaguchi, K., Kageyama, R. and Sano, M. (2017) ‘Topological defects control collective dynamics in neural progenitor cell cultures’, Nature. Nature Publishing Group, 545(7654), pp. 327–331. doi: 10.1038/nature22321.

Klingner, C. et al. (2014) ‘Isotropic actomyosin dynamics promote organization of the apical cell cortex in epithelial cells.’, The Journal of cell biology, 207(1), pp. 107–21. doi: 10.1083/jcb.201402037.

Lecuit, T. and Lenne, P.-F. P.-F. (2007) ‘Cell surface mechanics and the control of cell shape, tissue patterns and morphogenesis’, Nat Rev Mol Cell Biol. Nature Publishing Group, 8(8), pp. 633–644. Available at: http://dx.doi.org/10.1038/nrm2222.

Mark, S. et al. (2010) ‘Physical model of the dynamic instability in an expanding cell culture’, Biophysical Journal. Elsevier, 98(3), pp. 361–370. doi: 10.1016/j.bpj.2009.10.022.

Monier, B. et al. (2010) ‘An actomyosin-based barrier inhibits cell mixing at compartmental boundaries in Drosophila embryos.’, Nature cell biology. Nature Publishing Group, 12(1), pp. 60–65; sup pp 1-9. doi: 10.1038/ncb2005.

Mueller, R., Yeomans, J. M. and Doostmohammadi, A. (2019) ‘Emergence of Active Nematic Behavior in Monolayers of Isotropic Cells’, Physical Review Letters. American Physical Society, 122(4), p. 48004. doi: 10.1103/PhysRevLett.122.048004.

Nishimura, T. and Takeichi, M. (2008) ‘Shroom3-mediated recruitment of Rho kinases to the apical cell junctions regulates epithelial and neuroepithelial planar remodeling.’, Development (Cambridge, England), 135(8), pp. 1493–1502. doi: 10.1242/dev.019646.

Ostuni, E. et al. (2000) ‘Patterning Mammalian Cells Using Elastomeric Membranes’, Langmuir, 16(20), pp. 7811–7819.

Rauzi, M. et al. (2008) ‘Nature and anisotropy of cortical forces orienting Drosophila tissue morphogenesis.’, Nature cell biology, 10(12), pp. 1401–10. doi: 10.1038/ncb1798.

Rauzi, M., Lenne, P.-F. and Lecuit, T. (2010) ‘Planar polarized actomyosin contractile flows control epithelial junction remodelling.’, Nature. Nature Publishing Group, 468(7327), pp. 1110–4. doi: 10.1038/nature09566.

Reffay, M. et al. (2011) ‘Orientation and polarity in collectively migrating cell structures: statics and dynamics.’, Biophysical journal. Biophysical Society, 100(11), pp. 2566–75. doi: 10.1016/j.bpj.2011.04.047.

Reffay, M. et al. (2014) ‘Interplay of RhoA and mechanical forces in collective cell migration driven by leader cells.’, Nature cell biology, 16(3), pp. 217–23. doi: 10.1038/ncb2917.

Reinsch, S. and Karsenti, E. (1994) ‘Orientation of Spindle Axis and Distribution of Plasma Membrane Proteins during Cell Division in Polarized MDCKII Cells’, Journal of Cell Biology, 126(6), pp. 1509–1526.

Riveline, D. et al. (2001) ‘Focal contacts as mechanosensors: externally applied local mechanical force induces growth of focal contacts by an mDia1-dependent and rock-independent mechanism’, J Cell Biol, 153(6), pp. 1175–1186. doi: 10.1083/jcb.153.6.1175.

Saw, T. B. et al. (2017) ‘Topological defects in epithelia govern cell death and extrusion’, Nature. Nature Publishing Group, 544(7649), pp. 212–216. doi: 10.1038/nature21718.

Serra-Picamal, X. et al. (2012) ‘Mechanical waves during tissue expansion’, Nature Physics. Nature Publishing Group, 8(8), pp. 628–634. doi: 10.1038/nphys2355.

Shyer, A. E. et al. (2013) ‘Villification: How the Gut Gets Its Villi’, Science, 342(6155), pp. 212–218. doi: 10.1126/science.1238842.Villification.

Solon, J. et al. (2009) ‘Pulsed Forces Timed by a Ratchet-like Mechanism Drive Directed Tissue Movement during Dorsal Closure’, Cell, 137(7), pp. 1331–1342. doi: 10.1016/j.cell.2009.03.050.

Tarle, V. et al. (2015) ‘Modeling the finger instability in an expanding cell monolayer’, Integrative Biology (United Kingdom). Royal Society of Chemistry, 7(10), pp. 1218–1227. doi: 10.1039/c5ib00092k.

Theveneau, E. and Linker, C. (2017) ‘Leaders in collective migration: Are front cells really endowed with a particular set of skills?’, F1000Research, 6, pp. 1–11. doi: 10.12688/f1000research.11889.1.

Vishwakarma, M. et al. (2018) ‘Mechanical interactions among followers determine the emergence of leaders in migrating epithelial cell collectives’, Nature Communications. Springer US, 9(1). doi: 10.1038/s41467-018-05927-6.

Vuong-Brender, T. T. K., Yang, X. and Labouesse, M. (2016) ‘Chapter Thirty-Five - C. elegans Embryonic Morphogenesis’, in Wassarman, P. (ed.) Essays on Developmental Biology, Part A. Academic Press, pp. 597–616. doi: https://doi.org/10.1016/bs.ctdb.2015.11.012.

Zhang, H. et al. (2011) ‘A tension-induced mechanotransduction pathway promotes epithelial morphogenesis.’, Nature, 471(7336), pp. 99–103. doi: 10.1038/nature09765.

Zhang, H. and Labouesse, M. (2012) ‘Signalling through mechanical inputs - a coordinated process’, Journal of Cell Science, 125(13), pp. 3039–3049. doi: 10.1242/jcs.093666.

