## Supplementary Information for "Epithelial colonies *in vitro* elongate through collective effects"

#### Contents:

Figure Supplements

Supplementary Information

Movie Captions

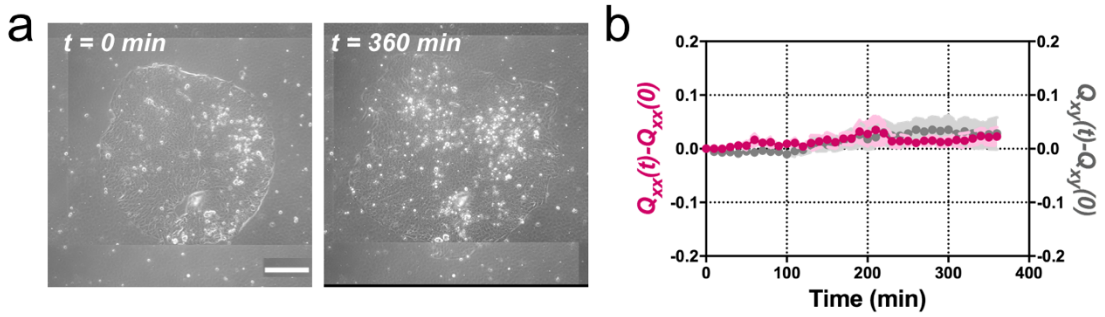


**Figure 1 – figure supplement 1.** ***MDCK colonies of 750 µm in diameter expand isotropically***. (**a**) Phase contrast images of the spontaneous expansion of a MDCK colony for 360 min. Scale bar 200 µm. (**b**) $Q_{xx}$ (left y axis) and $Q_{xy}$ (right y axis) during 360 min of colony expansion. Mean value ± standard error of the mean, n = 3 colonies from N = 3 independent experiments.

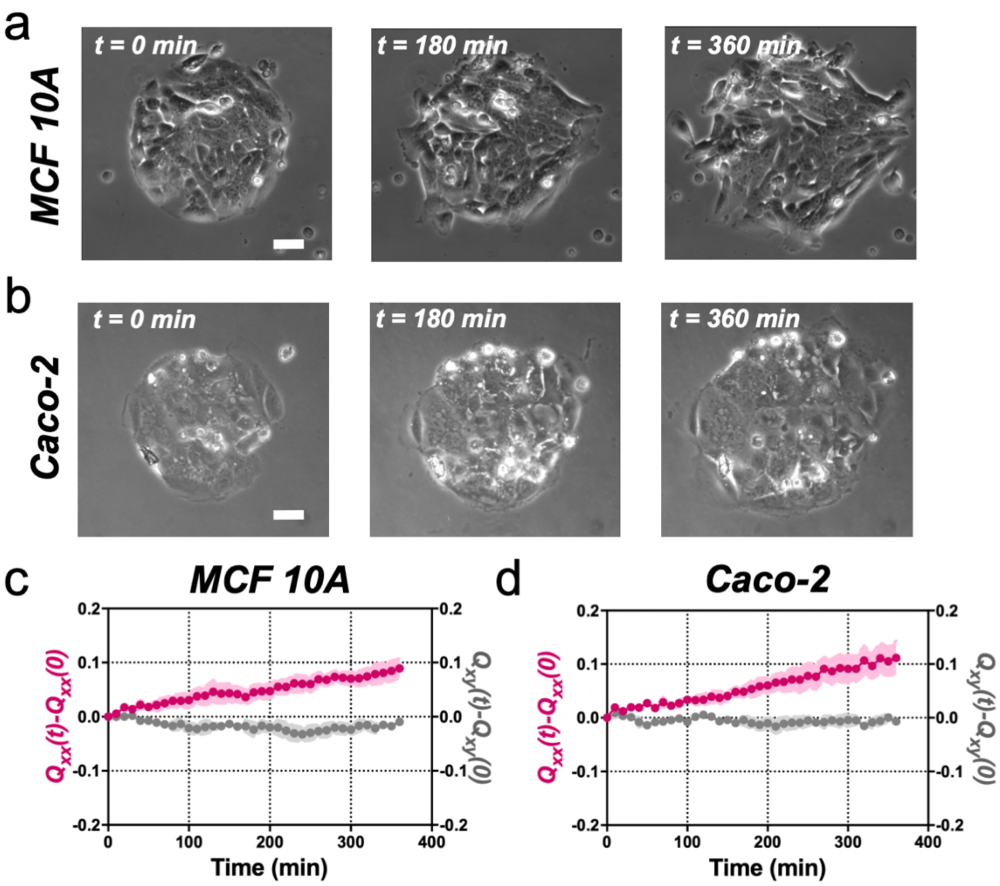


**Figure 1 – figure supplement 2.** ***MCF 10A and Caco-2 colonies of 250 µm in diameter expand asymmetrically similar to MDCK colonies***. (**a**) Phase contrast images of the spontaneous elongation of a MCF 10A colony for 360 min. Scale bar 50 µm. (**b**) Phase contrast images of the spontaneous elongation of a Caco-2 colony for 360 min. Scale bar 50 µm. (**c**) $Q_{xx}$ (left y axis) and $Q_{xy}$ (right y axis) during 360 min of MCF 1A colony expansion. Mean value ± standard error of the mean, n = 10 colonies from N = 3 independent experiments. (**d**) $Q_{xx}$ (left y axis) and $Q_{xy}$ (right y axis) during 360 min of Caco-2 colony expansion. Mean value ± standard error of the mean, n = 6 colonies from N = 3 independent experiments.

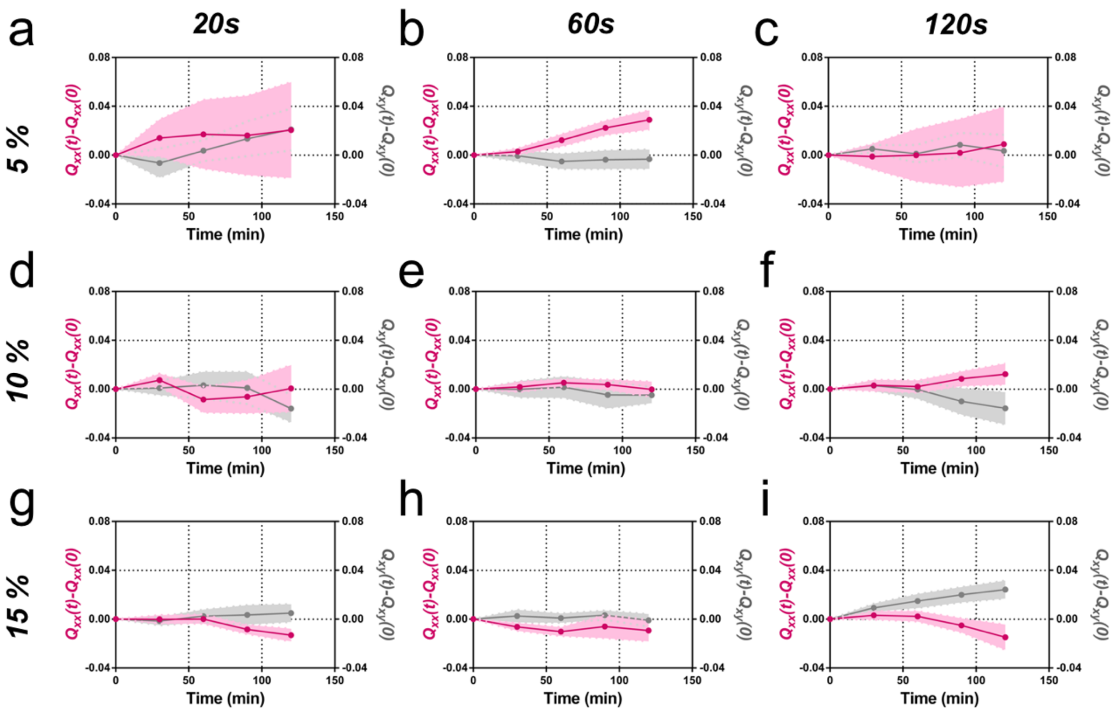


**Figure 2 – figure supplement 1.** ***Elongation as a function of frequency and amplitude of cyclic uniaxial stretching***. Cumulative $Q_{xx}$ (left y axis) and cumulative $Q_{xy}$ (right y axis) during 120 min of colony expansion ($\alpha = 0$). Mean value ± standard error of the mean. (**a**) 5% amplitude and 20s frequency N = 3, n = 5 colonies. (**b**) 5% amplitude and 60s frequency N = 9, n = 20 colonies. (**c**) 5% amplitude and 120s frequency N = 5, n = 6 colonies. (**d**) 10% amplitude and 20s frequency N = 4, n = 6 colonies. (**e**) 10% amplitude and 60s frequency N = 5, n = 8 colonies. (**f**) 10% amplitude and 120s frequency N = 4, n = 8 colonies. (**g**) 15% amplitude and 20s frequency N = 3, n = 9 colonies. (**h**) 15% amplitude and 60s frequency N = 3, n = 6 colonies. (**i**) 15% amplitude and 120s frequency N = 3, n = 6 colonies.

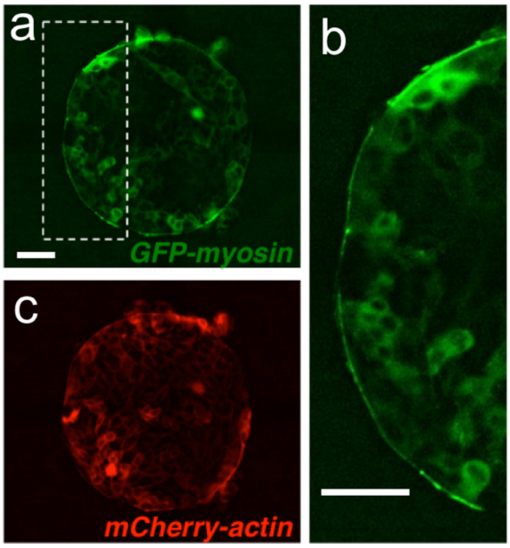


**Figure 4 – figure supplement 1.** ***A supra-cellular acto-myosin cable sets the boundary of the colony***. (**a**) MDCK cells expressing GFP-myosin and mCherry-actin. (**b**) Inset of the myosin cable at the colony boundary. (c) Image of the actin signal. Scale bar 50 µm.

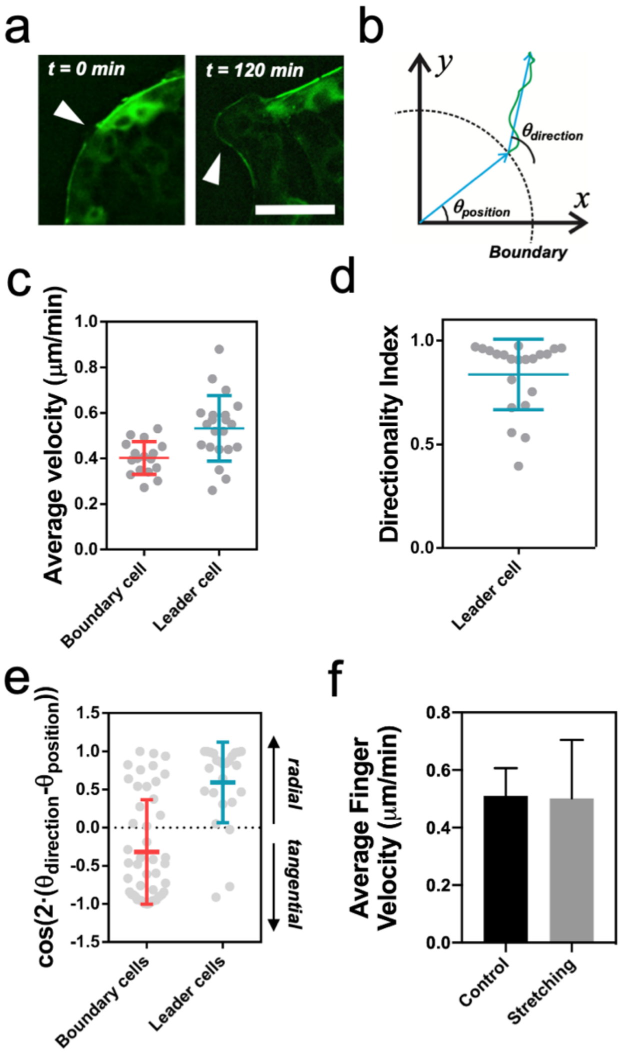


**Figure 4 – figure supplement 2.** ***Leader cell dynamics***. (**a**) Discontinuities in the myosin cable structure at t = 0 min (white arrows) become finger-like structures at t = 120 min (white arrows). Scale bar 50 µm. (**b**) Schematics of a cell trajectory. Its initial position at the boundary with respect to the center of the colony defines the angle $\theta_{position}$. The vector between the initial and the final position of the cell defines the trajectory direction $\theta_{direction}$. (**c**) Average velocity of cells at the outer rim of the colony. Leader cells are faster than other boundary cells. $n_{boundary}$ = 17 cells from 8 colonies and N = 4 independent experiments and $n_{leader}$ = 21 cells from 12 colonies and N = 5 independent experiments. Individual cells in grey with Mean ± SD. (**d**) Directionality index for leader cells as the ratio of the distance between the initial and final position over the total distance covered by the cell. $n_{leader}$ = 21 cells from 12 colonies and N = 5 independent experiments. Individual cells in grey with Mean ± SD. (**e**) Cosine of two times the angle difference between $\theta_{position}$ and $\theta_{direction}$ for cells at the boundary and cells in finger-like structures. Values close to 1 correspond to radial trajectories and values close to -1 correspond to tangential trajectories. Mean value ± standard deviation, $n_{boundary}$ = 49 cells from 8 colonies and N = 4 independent experiments and $n_{leader}$ = 25 cells from 12 colonies and N = 5 independent experiments. Individual cells in grey with Mean ± SD. (**f**) Average finger velocity in control and in stretching cases. Only fingers parallel to the direction of stretching ($\Delta\theta < 30º$) from Figure 4d are considered. Mean ± SD.

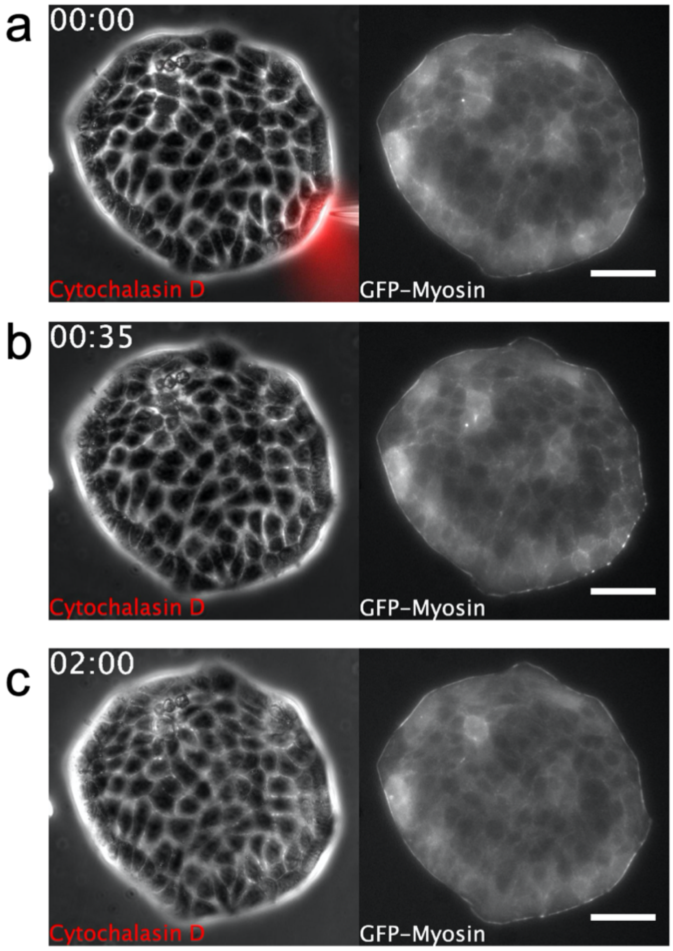


**Figure 5 – figure supplement 1. S*upra-cellular acto-myosin cable disruption***. Phase contrast images of a MDCK colony expressing GFP-myosin. A micropipette is placed shortly next to the colony boundary in (a). Note dynamics of myosin cable at the boundary. Scale bar 50 µm. Time: hh:mm. (**a**) Cytochalasin D is locally injected. (**b**) Disassembly of the myosin cable. (**c**) Recovery after Cytochalasin D injection. The myosin cable reforms at the colony boundary and no growth of finger is observed (see also the full dynamics in Movie 5).

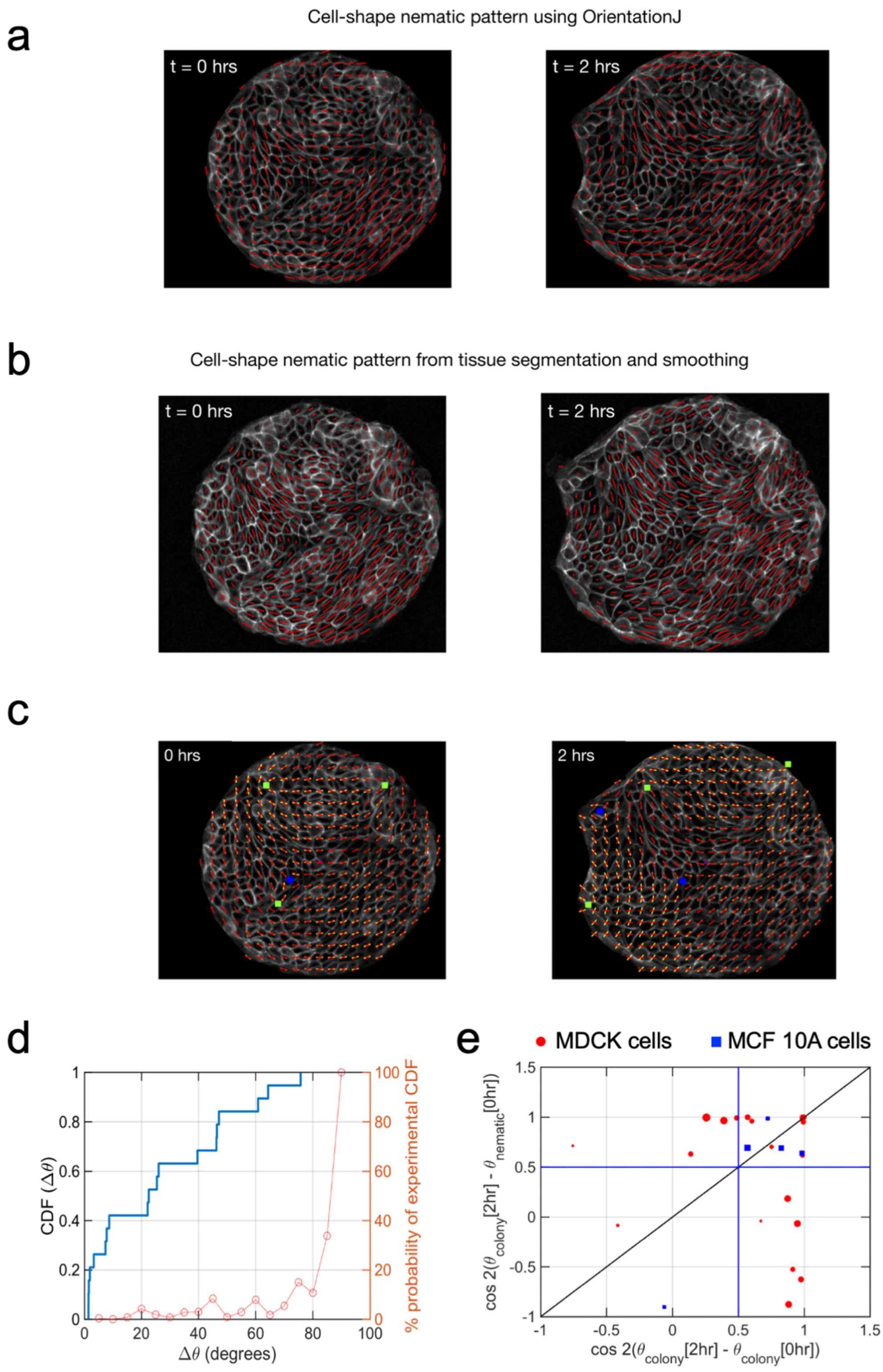


**Figure 5 – figure supplement 2.** ***Cell shape nematic field.*** Comparison of (**a**) cell shape nematic field obtained from OrientationJ plugin of ImageJ with (**b**) its counterpart obtained from tissue segmentation, triangulation and smoothing. It can be seen that the simpler, direct approach in (**a**) shows a very similar output as from the more detailed procedure in (**b**). (**c**) Fitting of cell-shape nematic field (yellow segments) with the one obtained experimentally (red segments) from OrientationJ using a simple expression (Eq. 14 of Supplementary Material) in terms of topological defects. The green squares and the blue circles represent, respectively, +1/2 and -1/2 topological defects at the corresponding locations obtained from the experimental nematic field. (**d**)

Correlation between $\theta_{nematic}$ (0 hrs) and $\theta_{colony}$(2 hrs) for finite sample size. Simple calculation to obtain the probability of observing a cumulative distribution function (CDF) for $\Delta\theta$, the angle between the mean colony shape nematic $\theta_{nematic}$ at 0 hrs and overall colony elongation $\theta_{colony}$ at 2 hrs, in the absence of any physical correlation between these two quantities, i.e., $\Delta\theta$ is picked with uniform probability from the set [0°, 90°] (orange curve and y axis on the right). It can be seen that in such a case, the probability to experimentally observe CDF > 0.6 for $\Delta\theta = 30°$ in a sequence of 19 independent experiments is less than 1/100. This provides a strong indication that $\theta_{nematic}$(0 hrs) is very likely to be physically connected with $\theta_{colony}$(2 hrs) and not randomly picked between 0° and 90°. n = 19 colonies of N = 5 independent experiments. (**e**) Final orientation of the colony is correlated with the initial shape nematic of initial orientation. The orientation of mean cell shape nematic $\theta_{nematic}$ of the colony at 0 hrs is strongly correlated with the overall orientation of the colony $\theta_{colony}$ at 2 hrs. It can be inferred from the presence of 17 out of 26 colonies above the horizontal blue line at 0.5 (corresponding to $\Delta\theta = 30°$). Of the rest of the 9 colonies which do not show this trend, 7 colonies show a strong correlation of their final orientation with their orientation at the start of experiment. The size of the symbols is proportional to the magnitude of anisotropy of the colony shape after 2 h. $n_{MDCK} = 19$ colonies of $N_{MDCK} = 5$ independent experiments. $N_{MCF\ 10A} = 5$ colonies of $N_{MCF\ 10A} = 2$ independent experiments.

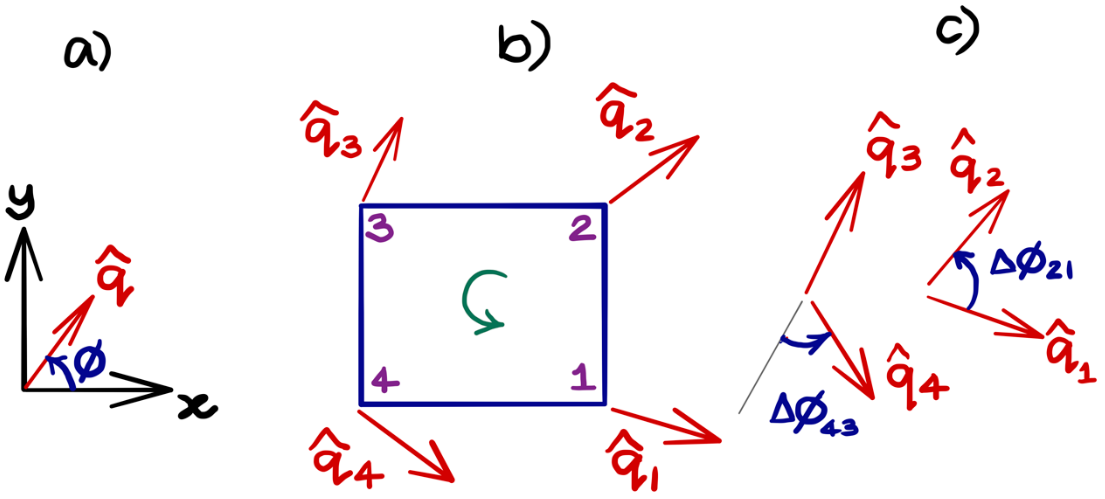


**Figure 5 – figure supplement 3: Obtaining topological charge for the orientation field $\hat{\boldsymbol{q}}$ on a rectangular grid.** (**a**) An orientation director $\hat{\boldsymbol{q}} = \cos\phi\, \hat{\boldsymbol{e}}_x + \sin\phi\, \hat{\boldsymbol{e}}_y$ makes an angle ϕ with the axis as shown. (**b**) The orientation field at the nodes of the smallest cell of the rectangular grid. The topological charge of $\hat{\boldsymbol{q}}$ within the cell is obtained by calculating the change in the orientation angle $\Delta\phi$ between the adjacent nodes of the grid and traversed in the anticlockwise direction. (**c**) The angle $\Delta\phi$ is taken to be the smallest of the angles between the adjacent directors and respects the nematic nature (or $\hat{\boldsymbol{q}}$ and $-\hat{\boldsymbol{q}}$ symmetry) of the field. We note that this procedure is ambiguous when $\Delta\phi$ is a multiple of π/2, a case that generally does not happen in practice. After using anticlockwise $\Delta\phi > 0$ and clockwise $\Delta\phi < 0$, the net topological charge of the orientation field $\hat{\boldsymbol{q}}$ within the rectangular cell in (b) is obtained as $k = \frac{1}{2\pi}\sum_j \Delta\phi_{j+1,j}$.

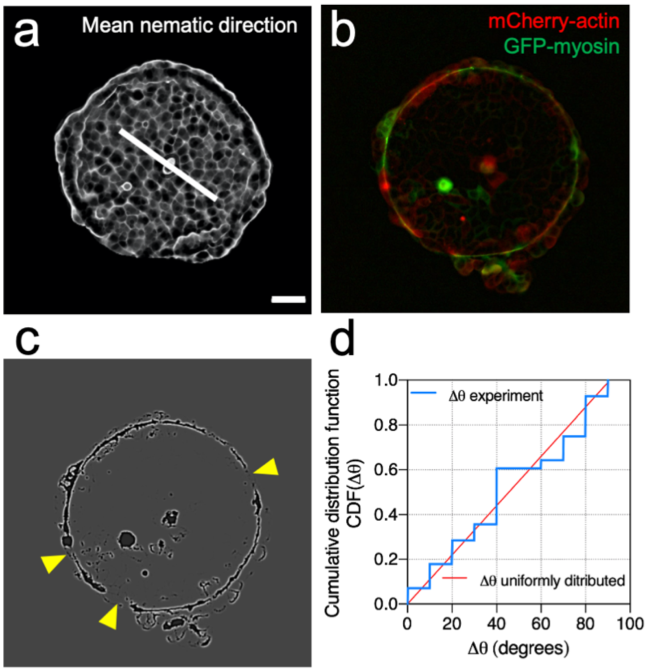


**Figure 5 – figure supplement 4:** ***Mean nematic direction and defects in the acto-myosin cable.*** (**a**) Mean cell shape nematic of a MDCK colony. Scale bar 50 µm. (**b**) Actin (red) and myosin (green) signals at the boundary of the colony. (**c**) Processed image to identify openings of the cable (yellow arrows). (**d**) Cumulative distribution function of Δθ. Where Δθ corresponds to the angular distance between the mean nematic direction and the position of defects in the acto-myosin cable, $\Delta\theta = \left|\theta_{nematic} - \theta_{defect}\right|$. N = 3 experiments, n = 7 colonies.

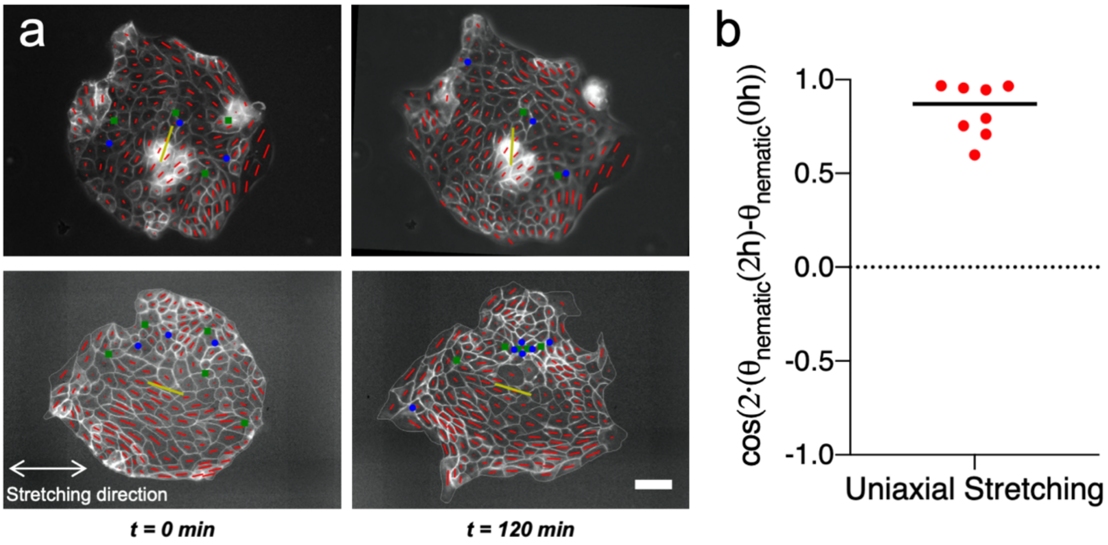


**Figure 5 – figure supplement 5:** ***Uniaxial cyclic stretching does not modify the mean nematic direction.*** (**a**) Cell shape nematic field (red segments) and mean nematic direction (yellow) of 2 colonies under uniaxial cyclic stretching at t = 0 min and t = 120 min. Mean nematic direction remains invariant after 2 hours of uniaxial cyclic stretching. Scale bar 50 µm. (**b**) Box plot of the change in the mean nematic direction of colonies under uniaxial cyclic stretching, $\cos(2\cdot(\theta_{nematic}(2\ h) - \theta_{nematic}(0\ h))$. n = 8 colonies from N = 6 experiments.

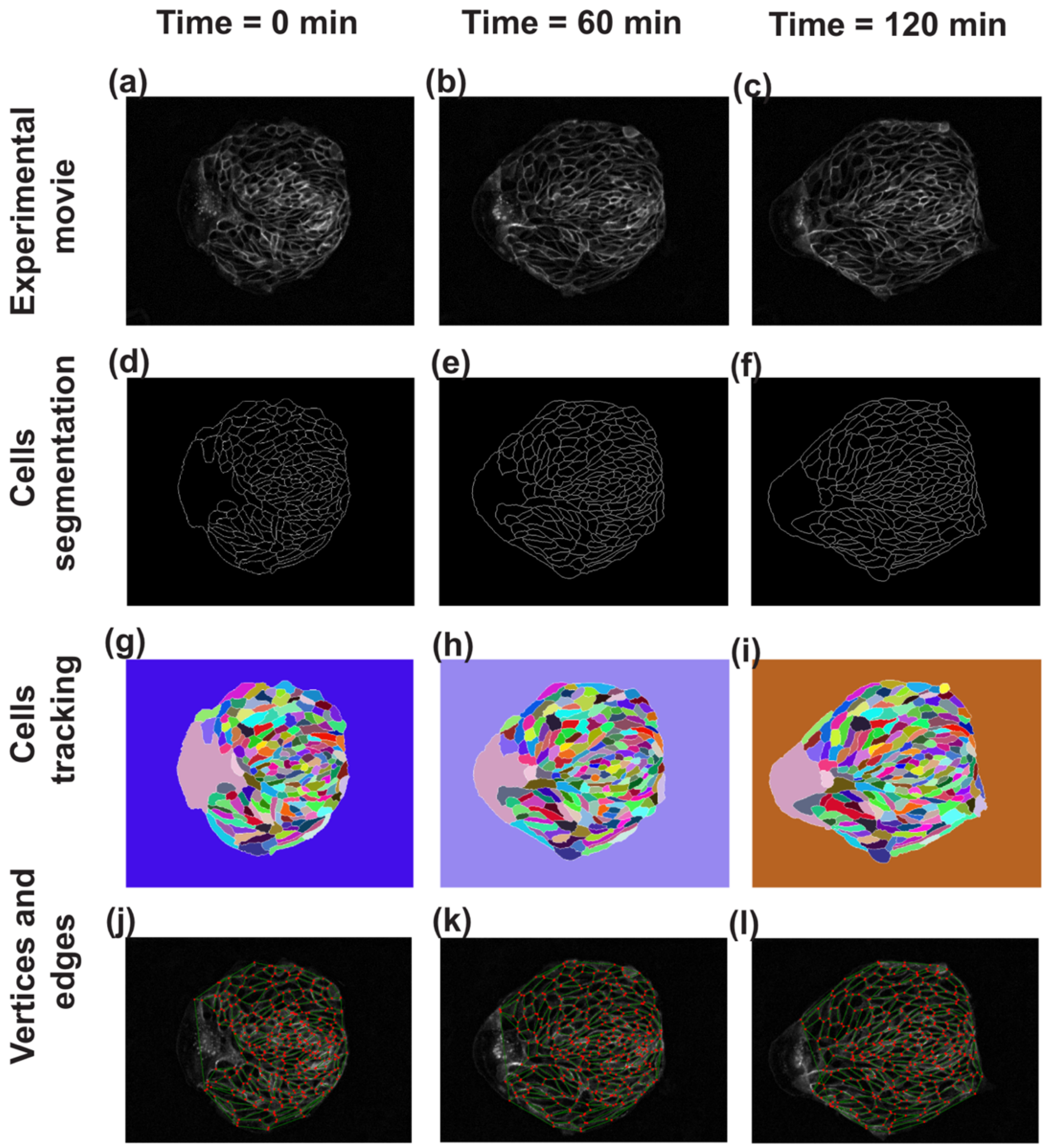


**Figure 6 – figure supplement 1.** ***Steps in image analysis.*** As the first step of analysis using TissueMiner, the images obtained from the experiment have been skeletonized, segmented and tracked using TissueAnalyser. (**a**–**c**) Show the images of cell colony at three different time points. (**d**–**f**) In order to differentiate the cell edges from the rest of the tissue for easy edge detection, the tissue has been skeletonized by adjusting the threshold value. (**g**–**i**) TissueAnalyser detects the edges and segments the colony, based on a watershed algorithm. At this step, algorithm also tracks the cells and assigns them a color and a global track identity, which remain constant throughout the process. When a cell undergoes division, a new track identity and color are assigned to one of the two daughter cells, while keeping the other same. (**j**–**l**) Using the output from TissueAnalyser, TissueMiner analyzes the tissue and represents it as a collection of cells made of a network of edges (green lines) connected to vertices (red dots) at any given time.

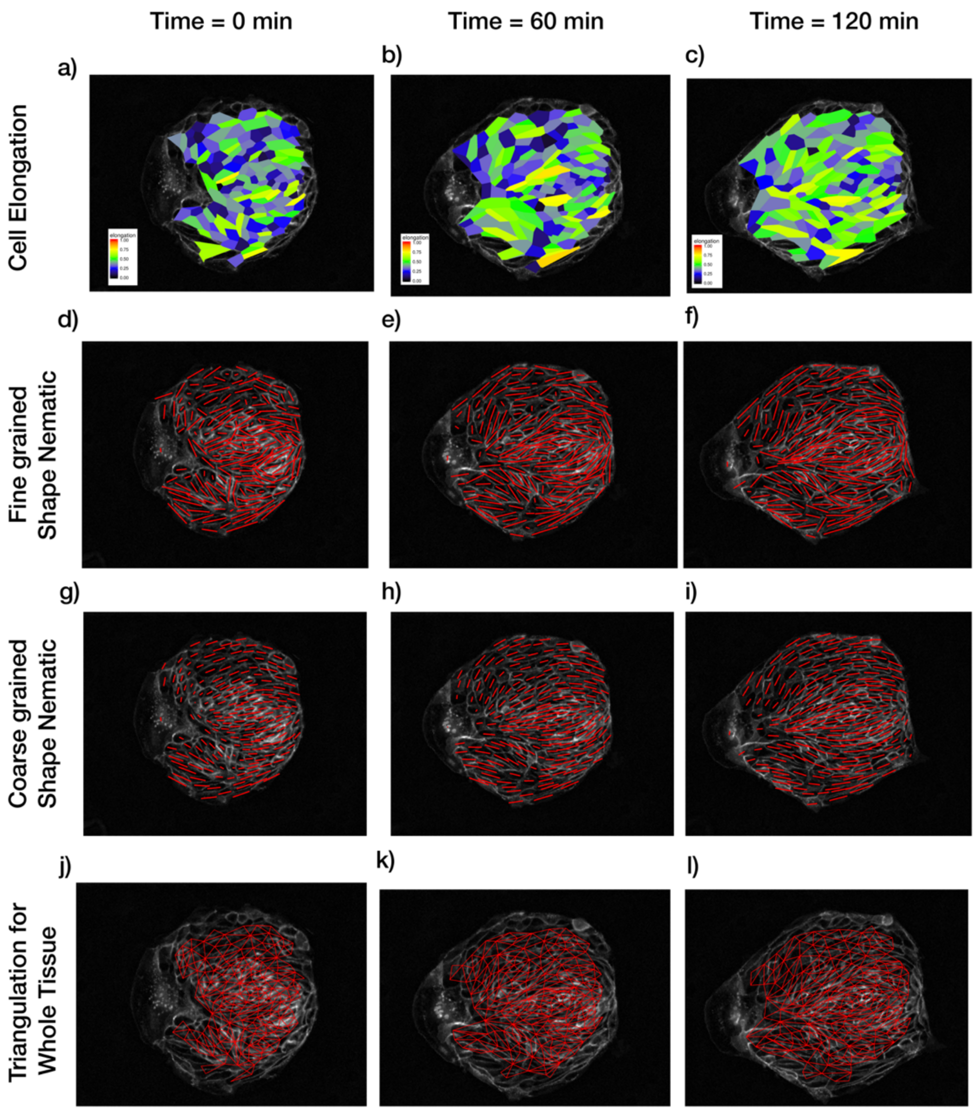


**Figure 6 – figure supplement 2.** ***Patterns of cellular parameters obtained from TissueMiner*** **analysis.** (**a** – **c**) Cell elongation magnitude distribution is plotted for three different time points. (**d** – **f**) Actual fine-grained cell elongation nematic for individual cells. The length of the segments is proportional to the magnitude of cell-shape nematic $\varepsilon$ and the orientation is $\varphi$ (see Supplementary Material Section V) (**g** – **i**) Coarse-grained cell shape nematic is obtained by using a Gaussian kernel of size 30 pixels around every cell. (**j** – **l**) The algorithm to find out tissue deformation is based on the triangulation method as described in Refs. [23, 27] of the main paper. Any topological changes in the form of T1 transition, cell division or cell extrusion will lead to a new triangulation. The triangulation is not reliable near the boundary and hence a layer of cells near the boundary is not included in the triangulation and subsequent calculations for tissue pure shear.

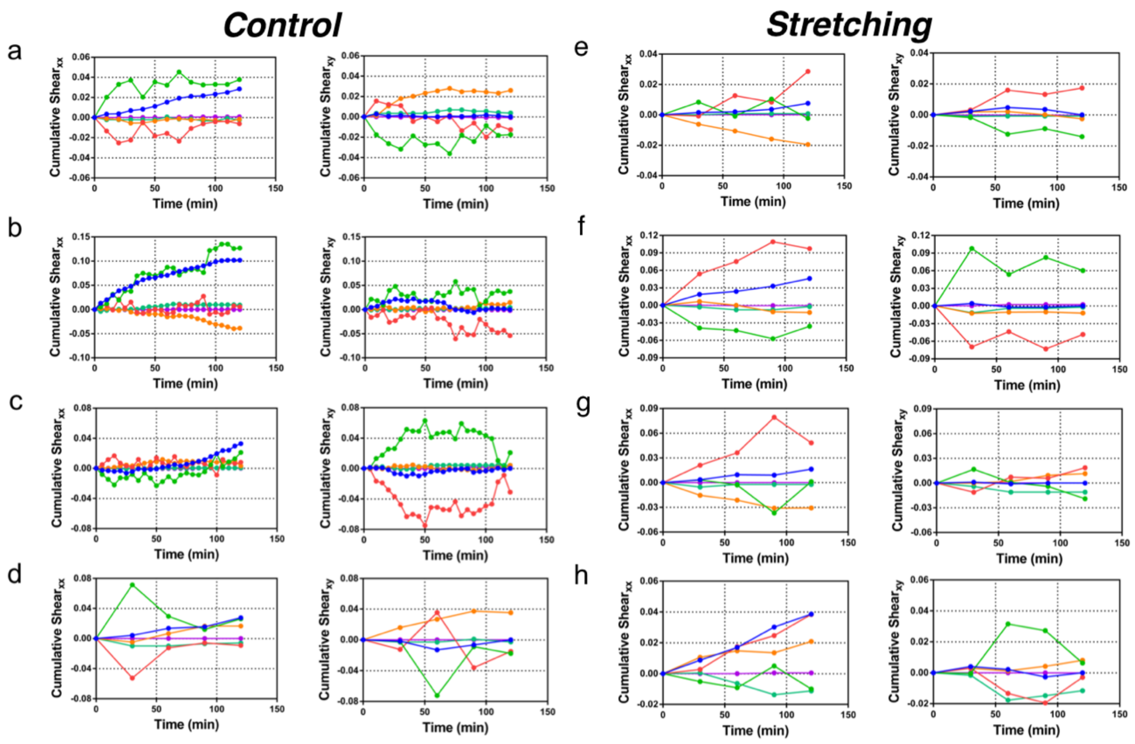


**Figure 6 – figure supplement 3.** ***Cumulative pure shear decomposition.*** (**a-d**) Control experiments. Decomposition of pure shear (blue) into its components arising from cell shape change (green), T1 transitions (red), correlation effects (orange), cell division (turquoise), and cell extrusion (purple). The *x* direction for each colony is chosen such that *xy* component of total shear is zero at t = 120 mins, and indicates the overall orientation of colony anisotropy. The anti-correlation effect between cell elongation and other components that is seen in Fig. 4f of the main paper is clearly visible in every case. The interval between subsequent time data points is 10 min for (**a**), 5 mins in (**b-c**) and 30 mins for (**d**). The data for *xx* component of pure shear in (**a**) is presented in Figure 6. (**e-h**) Stretched colonies. Same analysis method and time evolution of pure shear presented in (**a-d**) for control colonies, but now for stretched colonies. The time interval between subsequent time data points is 30 mins. Interestingly, the anti-correlation effect between the cell elongation and other components with respect to the total pure shear is seen here too.

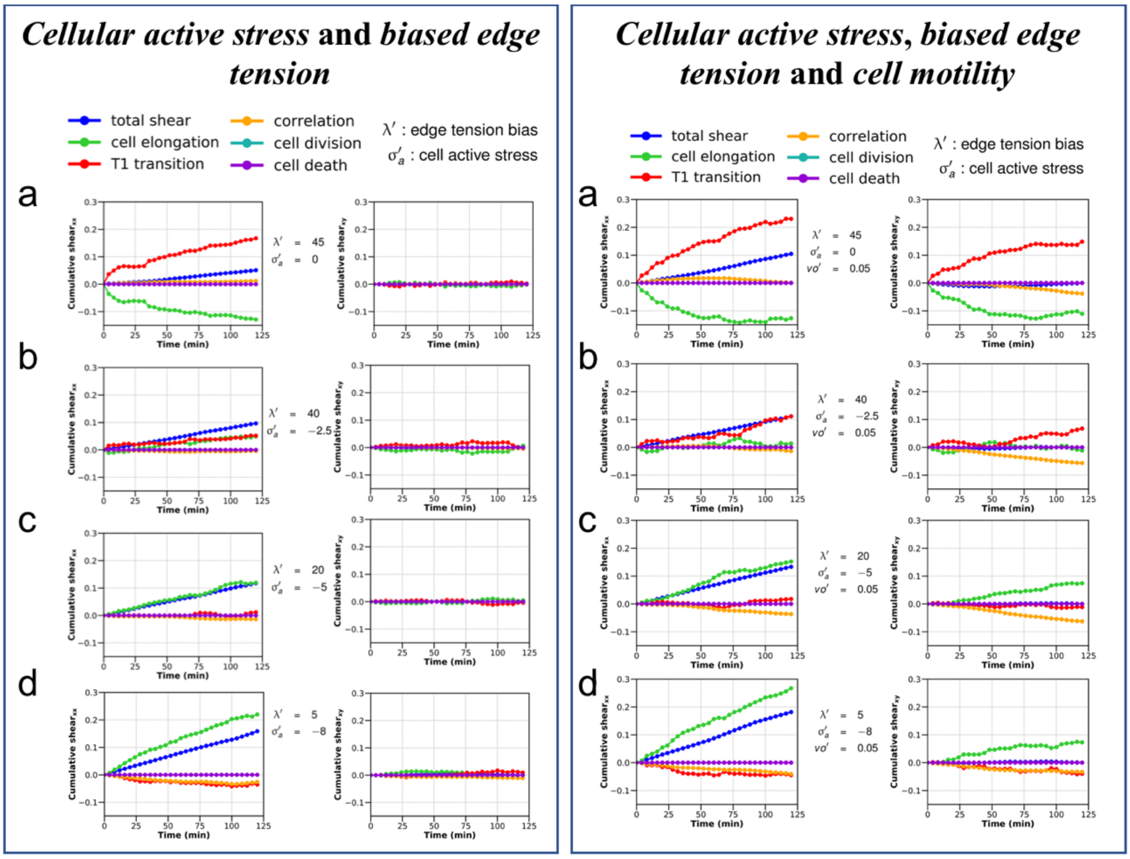


**Figure 7 – figure supplement 1.** ***Cumulative pure shear decomposition patterns for the simulated colonies.*** (**Left**) Varying strength of cellular active stress and biased edge tension. Briefly, the *cellular active stress* $(\sigma_a')$ is an internal cellular stress that tends to drive cell elongation. Biased edge tension $(\lambda')$ is implemented by modulating the line tension of the cell edges on the basis of the relative angle the edges makes with the polarity of their two shared cells. A colored noise with a small strength and persistence time is applied on the edge line tension to ensure baseline fluctuations. Note the relative positions of the pure shear due to T1 transitions (red) and cell elongations (green) with respect to the overall colony shear (blue). The *x* axis is chosen to be the major axis of elongation such that the *xy* component of cumulative total pure shear strain is zero at the end of the simulation. It can be seen from a-d that the relative position of the red and green curves with respect to the blue curve is an indication of the internal processes which are more dominant in a particular colony and show marked similarity with the corresponding experimental curves seen in Figure 6 – figure supplement 1. (**Right**) Varying strength of *cellular active stress*, *biased edge tension* and *cell motility.* All simulations shown here are as in left panel a-d, but with the addition of a small motility term $v_0'$. It can be clearly seen that addition of motility does not qualitatively modify the relative positions of the contributions from cell elongation (green) and T1 transitions (red) with respect to each other when compared with their counterparts in left panel. However, in this particular case, for the parameters and the initial conditions, the *xy* component of the shear is comparatively more dominant.

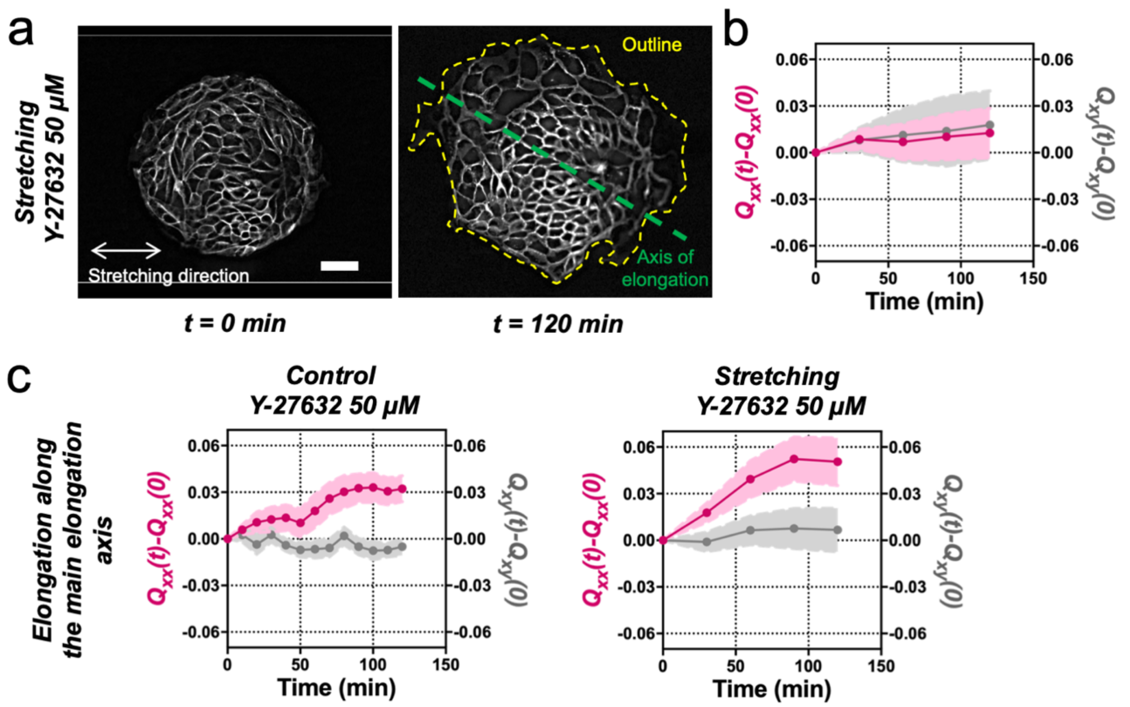


**Figure 8 – figure supplement 1.** ***Stretching-dependent elongation is mediated by ROCK.*** (**a**) Fluorescent images of an Ecadherin-GFP MDCK colony evolving for 120 min under cyclic mechanical stretching when ROCK is inhibited by Y-27632 50µM. The direction of the uniaxial stretching is indicated in white, the outline of the colony at 120 min is indicated in yellow and the major axis ($\theta(t_{final})$) is indicated in green. Colonies do not elongate along the stretching direction, but they still have a main elongation axis. Scale bar 50µm. (**b**) $Q_{xx}$ (left y axis) and $Q_{xy}$ (right y axis) of MDCK colonies during 120 min of cyclic uniaxial stretching in the presence of ROCK inhibitor. Elongation is measured in the direction of external force ($\alpha = 0$). Mean value ± standard error of the mean. N = 3, n = 8 colonies. (**c**) Comparison of the cumulative $Q_{xx}$ (left y axis) and cumulative $Q_{xy}$ (right y axis) during 120 min of colony expansion when ROCK is inhibited in control and under cyclic uniaxial stretching. Elongation is measured along the spontaneous elongation direction ($\alpha = \theta(t_{final})$). Mean value ± standard error of the mean, $N_{control} = 3$, n = 12 colonies and $N_{stretching} = 3$, n = 8 colonies.

### Supplementary Information: Comelles et al.

#### I. QUANTIFICATION OF TISSUE KINEMATICS

In the first step of analysis, time-lapse images of MDCK colonies are segmented to partition the tissue into individual cells and then tracked for their movements using Tissue-Analyser (TA) plugin of FIJI [1, 2] (see Figure 6 – figure supplement 1a-i). This information of the tracked cell colony is then passed to Tissue-Miner (TM), which extracts data and stores the cellular geometry and connectivity in the form of cell vertices and bonds using a *snakemake* based automated workflow (see Figure 6 – figure supplement 1j-l). As described in Refs. [3, 4], TM has *R* or *Python* API that can be used to quantify fine-grained (cell-level) and coarse-grained (colony-level) deformation properties of epithelial colonies. Even though the scripts to calculate the basic kinematic quantities are available, users have the freedom to add new computational routines to calculate the quantities of their interest.

##### A. Analysis of cell elongation

Cell-shape anisotropy is an important marker of stress in the cells in response to either internal activity or external forcing [5, 6]. To understand and quantify the elongation of cells over time, TM computes an elongation parameter proposed in Ref. [1]. In their study to understand the role of flow in orienting the axis of polarity in *Drosophila* wings, Aigouy et al. [1] use a nematic tensor with a magnitude and orientation to quantify the elongation of each cell. For a given cell with area $A_c$, the elongation nematic tensor is proposed as

$$E = \begin{bmatrix} \epsilon_{xx} & \epsilon_{xy} \\ \epsilon_{xy} & -\epsilon_{xx} \end{bmatrix}. \tag{1}$$

This tensor $E$ is traceless and symmetric with components given by

$$\begin{aligned} \epsilon_{xx} &= \frac{1}{A_c}\int_0^{2\pi}\int_0^{R(\phi)} \cos(2\phi) r dr d\phi \\ \epsilon_{xy} &= \frac{1}{A_c}\int_0^{2\pi}\int_0^{R(\phi)} \sin(2\phi) r dr d\phi \end{aligned}, \tag{2}$$

where $R(\phi)$ is the distance to cell boundary at an angle $\phi$ (SI of Ref. [1]). For this nematic tensor, the magnitude is given by

$$\epsilon = ({\epsilon_{xx}}^2 + {\epsilon_{xy}}^2)^{\frac{1}{2}}, \tag{3}$$

and the orientation is given by angle $\phi$

$$\begin{aligned} \cos(2\phi) &= \frac{\epsilon_{xx}}{\epsilon}, \\ \sin(2\phi) &= \frac{\epsilon_{xy}}{\epsilon}. \end{aligned} \tag{4}$$

Based on the above expressions, the spatial distribution of the magnitude of shape nematic for individual cells within the colony for three time points is shown in Figure 6 – figure supplement 2a-c. The fine-grained nematic segments for individual cells whose magnitude is proportional to $\epsilon$ and orientation is $\phi$ (Figure 6 – figure supplement 2d-f). The coarse-grained representation of this tensor (Figure 6 – figure supplement 2g-i) is obtained by spatially averaging the elongation nematic over several cells by using a Gaussian smoothing kernel of size comparable to the size of cells (30 pixels in this case). From the figures it is clear the elongation of interior cells is strongly correlated with the orientation of the future 'finger' even at $t = 0$ h.

##### B. Analysis of tissue deformation from components of pure shear strain via triangulation

As discussed earlier, during different stages of morphogenesis, epithelial tissues are reported to undergo a series of physical changes. The various internal and external forces that are being applied to the tissue during these processes result in deformation and rearrangement of constituent cells. The shape modification rate of the tissue is quantified in terms of the anisotropic component of shear strain rate. However, at the tissue level in epithelial sheets, shear strain can arise due to a combination of cell elongation, T1 transitions, cell division, cell extrusion, and correlation effects. By studying each both component separately and with respect to the other components, one can get hints regarding the mechano-chemical state of the colony. As is described in Ref. [4], for any general 2-D velocity vector $\mathbf{v}$, the gradient tensor can be defined as

$$v_{ij} = \partial_i v_j = \begin{bmatrix} \frac{\partial v_x}{\partial x} & \frac{\partial v_y}{\partial x} \\ \frac{\partial v_x}{\partial y} & \frac{\partial v_y}{\partial y} \end{bmatrix}. \quad (5)$$

The above velocity gradient tensor for a group of cells can be split into three parts: (i) isotropic component to obtain the rate of area change, (ii) symmetric traceless component to quantify the pure shear strain rate, and (iii) an antisymmetric part that represents the local vorticity [4, 7] as follows

$$\begin{aligned} v_{ij} &= \frac{1}{2} v_{kk}\delta_{ij} + \tilde{v}_{ij} + \omega\delta_{ij} \\ &= \underbrace{\frac{1}{2}\begin{bmatrix} \frac{\partial v_x}{\partial x} + \frac{\partial v_y}{\partial y} & 0 \\ 0 & \frac{\partial v_x}{\partial x} + \frac{\partial v_y}{\partial y} \end{bmatrix}}_{v_{kk}\text{:isotropic}} + \underbrace{\frac{1}{2}\begin{bmatrix} \frac{\partial v_x}{\partial x} - \frac{\partial v_y}{\partial y} & \frac{\partial v_y}{\partial x} + \frac{\partial v_x}{\partial y} \\ \frac{\partial v_y}{\partial x} + \frac{\partial v_x}{\partial y} & \frac{\partial v_y}{\partial y} - \frac{\partial v_x}{\partial x} \end{bmatrix}}_{\tilde{v}_{ij}\text{:symmetric, anisotropic}} + \underbrace{\frac{1}{2}\begin{bmatrix} 0 & \frac{\partial v_y}{\partial x} - \frac{\partial v_x}{\partial y} \\ \frac{\partial v_x}{\partial y} - \frac{\partial v_y}{\partial x} & 0 \end{bmatrix}}_{\omega\text{:antisymmetric}}. \end{aligned} \quad (6)$$

Here, more specifically, $v_{kk} = v_{xx} + v_{yy}$ is the trace of the velocity gradient tensor, or the divergence of the velocity field and quantifies the rate of contraction or expansion in the cell area. The traceless symmetric part $\tilde{v}_{ij}$ is the pure shear strain rate and quantifies shape changes in the tissue. The antisymmetric component $\omega$ is the vorticity of the velocity field and provides values of rotational rates of the group of cells during deformation.

The total pure shear strain rate of the tissue $\tilde{v}_{ij}$, now represented in a tensorial form $\tilde{\mathbf{V}}$, can be further split into traceless, symmetric, components as follows

$$\tilde{\mathbf{V}} = \frac{D\mathbf{Q}}{Dt} + \mathbf{T} + \mathbf{C} + \mathbf{E} + \mathbf{D}, \quad (7)$$

where $\mathbf{Q}$ is the shear contribution coming from average cell elongation, $\mathbf{T}$ is the shear contribution from T1 transitions, $\mathbf{C}$ from cell divisions, $\mathbf{E}$ from cell extrusions and $\mathbf{D}$ due to the correlations effects in the tissue which arise due to coarse graining effects. Detailed derivation and discussion of this expression are available in Refs. [3, 4, 7]. The contributions from above mentioned components are quantified using the triangulation approach [3]. As shown in Figure 6 – figure supplement 2j-l, the triangulation method considers the entire colony without the boundary cells as a network of triangles, connecting the centers of the neighboring cells. Any topological changes in the form of cell division or extrusion will lead to a new triangulation. The shear strain rates in Eq. 7 can be integrated over time to provide cumulative shape deformation and the corresponding cumulative shear decomposition for the cell colony, and is obtained for the complete colony without including the external boundary cells (Figure 6b). Figure 4 of Ref. [7] schematically represents this decomposition of total tissue shear into individual cellular events.

In *Drosophila* pupal wing the $x$ axis is naturally provided by the A-P axis. In our *in vitro* situation, however, the $x$ axis is chosen as the dominant direction of colony elongation, such that at the end of the experiment ($t = 2$ h) the cumulative value of the total shear strain rate $\tilde{v}_{xy} = 0$. The cumulative shear decomposition plots for the experiments and simulations are shown in Figure 6b and Figure 6 – figure supplement 3 and Figure 7 – figure supplement 1.

##### C. Correlation between the orientation of mean cell-shape nematic and the orientation of the overall colony elongation

As discussed earlier, the shape of a cell can be thought of to be represented by a nematic tensor $\mathbf{Q}^{\text{cell}}$. Upon averaging this quantity over multiple cells, we obtain nematic field $\mathbf{Q}^{\text{nem}}(x, y, t)$ over the colony. As described in the previous sub-section, this nematic tensor is traceless and symmetric with an orientation $\phi(x, y, t)$ (Figure 5 – figure supplement 1 and Figure 6 – figure supplement 2g-i) and magnitude $Q^{\text{nem}}(x, y, t)$ such that

$$Q^{\text{nem}}_{xx}(x, y, t) = Q^{\text{nem}} \cos 2\phi \text{ and} \quad (8)$$
$$Q^{\text{nem}}_{xy}(x, y, t) = Q^{\text{nem}} \sin 2\phi. \quad (9)$$

This nematic field is obtained for our MDCK colonies by analyzing time-lapse images with the OrientationJ plugin of ImageJ [8, 9]. We verified that this image based approach gives results that are comparable with those obtained by spatially smoothing (with Gaussian kernel) the individual cell-shape nematics extracted from the segmented images of the colonies (Supplementary Material Section IA, Figure 5 – figure supplement 1 and Figure 6 – figure supplement 2). At any particular time $t$, the mean nematic field for the colony is

$$Q^{\mathrm{ncol}}_{xx}(t) = \langle Q^{\mathrm{nem}}_{xx}(x,y,t)\rangle_{x,y} \text{ and} \tag{10}$$

$$Q^{\mathrm{ncol}}_{xy}(t) = \langle Q^{\mathrm{nem}}_{xy}(x,y,t)\rangle_{x,y}. \tag{11}$$

The orientation of the mean cell-shape nematic is then simply

$$\theta_{\mathrm{nematic}}(t) = \frac{1}{2}\tan^{-1}\left(\frac{Q^{\mathrm{ncol}}_{xy}(t)}{Q^{\mathrm{ncol}}_{xx}(t)}\right), \tag{12}$$

where $\tan^{-1}$ is implemented in Matlab using the inbuilt function `atan2` [10]. In this analysis, the *coherence* of the image structure tensor is the counterpart of $Q^{\mathrm{col}}$, the magnitude of the mean cell-shape nematic [8]. We then compare the final colony orientation $\theta_{\mathrm{colony}}(t = 2 \text{ hrs})$, which is obtained from ellipse fitting, with the orientation of the mean colony nematic $\theta_{\mathrm{nematic}}(t = 0 \text{ hrs})$ (Figure 5b and Figure 5 – figure supplement 1). Also refer to Materials and Methods: Orientation field of the cells and topological defects in the Main Paper for additional details.

##### D. Topological defects as readouts of cell-shape nematic orientation field

Any nematic field can contain topological defects [11]. A visual inspection of the cell-shape orientation field $\mathbf{q} = \cos\phi\hat{\mathbf{e}}_x + \sin\phi\hat{\mathbf{e}}_y$ for MDCK colonies indicates the possible presence of topological defects (Figure 5 and Figure 5 – figure supplement 1). The strength $k$ of topological charge for this field (Supplementary Section IC) in a given region is obtained as

$$k = \frac{1}{2\pi}\oint \frac{d\phi}{ds}ds, \tag{13}$$

where the line integral is calculated by traversing the curve that encloses this region in an anticlockwise sense. For the experimental colonies, the nematic field is not continuous but obtained on the vertices of the grid-cell as shown in Figure 5 – figure supplement 2. In this case, the topological charge $k$ enclosed within the grid-cell is

$$k = \frac{1}{2\pi}\sum_j \Delta\Phi_{j+1,j}, \tag{14}$$

where the notation is as shown in Figure 5 – figure supplement 2. If no topological defect is present within the cell then $k = 0$. Else, the net topological charge would evaluate to $k = \pm 1/2$, corresponding to the effective strength of the topological defects within the cell.

The location and strength of topological defects thus obtained for the experimental data could then be used to approximately depict the entire nematic field as follows. We first model the energetics of the nematic orientation field with one constant Frank free energy [12]. If we then assume that the orientation field $\phi$ is in local mechanical equilibrium at every time-step, it will then satisfy the Laplace equation $\nabla^2\phi = 0$ [11]. The resulting orientation field $\phi(x, y)$ for the nematic field in the presence of a single topological defect of strength $k$ at the origin has a singular solution of the form

$$\phi = A\theta + \alpha, \tag{15}$$

where $\theta$ is the polar angle and $\alpha$ is a harmonic function that satisfies that Laplace equation [11]. However, for the current paper we find that taking $\alpha$ to be a constant that determines the orientation of the topological defect is sufficient to approximate the orientation nematic field (Figure 5d). Hence, as per Eqs. 13 and 15 the integral

$$\oint d\phi = 2\pi k, \tag{16}$$

for a closed curved around the defect gives $A = k$. If multiple defects are present in the domain, this solution for $\phi$ can be generalised to give

$$\phi_\alpha(x, y) = \alpha + \sum_i k_i \underbrace{\tan^{-1}\left(\frac{y - y_i}{x - x_i}\right)}_{\theta \text{ for defect } i}, \tag{17}$$

where $\{x_i, y_i\}$ and $k_i$ are the coordinates and the strength, respectively, of the $i^{\text{th}}$ nematic topological defect obtained from the experimental data, and $\alpha$ dictates the orientation of the defect. Here, the subscript $\alpha$ in $\phi_\alpha(x, y)$ is added to distinguish it from the experimentally obtained $\phi(x, y)$. As before, $\tan^{-1}$ is implemented in Matlab using the inbuilt function `atan2` [10]. We now use the fitting parameter $\alpha$ in $\phi_\alpha(x, y)$ to see how well it represents the orientation field $\phi(x, y)$ by using the following procedure. The experimental field is obtained on a grid with $N$ points $\{x_k, y_k\}$, where $k = 1, \ldots, N$. The difference between $\phi$ and $\phi_\alpha$ is quantified by the residue

$$r_\alpha = -\frac{1}{N}\sum_{k=1}^{N}\cos^2(\phi^k - \phi^k_\alpha) \tag{18}$$

obtained over the grid points $k$. The minimization of the value of $r_\alpha$ over $\alpha$ would provide us with a field where $\phi \approx \phi_\alpha$ in a mean sense. Interestingly, by using this simple approach with just one fitting parameter $\alpha$, we can get an excellent representation of the experimentally obtain nematic orientation field $\hat{\mathbf{q}}$ (Figure 5d).

##### E. Statistical test for cumulative distribution of the orientation differences between $\theta_{\text{nematic}}(0\text{ hrs})$ and $\theta_{\text{colony}}(2\text{ hrs})$

To check if the orientation of the colony elongation at $t = 2$ hrs is correlated with the orientation of the mean cell-shape nematic at $t = 0$ hrs, we examine the difference $\Delta\theta = |\theta_{\text{colony}}(2\,\text{hrs}) - \theta_{\text{nematic}}(0\,\text{hrs})|$. Specifically, as shown in Figure 4c, we obtain the cumulative distribution function (CDF) for $\Delta\theta$. If the quantity $\Delta\theta$ is completely random, i.e., $\theta_{\text{colony}}(2\,\text{hrs})$ is completely uncorrelated with $\theta_{\text{nematic}}(0\,\text{hrs})$, then the probability density for $\Delta\theta \in [0°, 90°]$ is equal to $1/90°$. In such a case the CDF($\Delta\theta$) as a function of $\Delta\theta$ should simply be a straight line with a slope of $1/90°$. However, it can clearly be seen that the experimentally observed CDF is significantly above this line, especially at relatively smaller values of $\Delta\theta \leq 30°$. This indicates that there indeed is correlation between $\theta_{\text{colony}}(2\,\text{hrs})$ and $\theta_{\text{nematic}}(0\,\text{hrs})$.

We now note that the total number of experiments (colonies) that were analysed for Figure 4c is $N = 19$. To check if the correlation between $\theta_{\text{colony}}(2\,\text{hrs})$ and $\theta_{\text{nematic}}(0\,\text{hrs})$ holds up for this finite sample size, we do the following. Let us take the probability density $p(\Delta\theta) = 1/90°$, $\forall \Delta\theta \in [0°, 90°]$, i.e., this relative angle $\Delta\theta$ is uniformly random between 0° and 90°. If we now sample $N = 19$ values for $\Delta\theta$ then we have created a *random* counterpart to the actual experimental result. We repeat this trial using a straightforward Matlab script and obtain $M = 10^6$ samples, each with $N = 19$ experiments. Of all the samples thus randomly generated, for a given $\Delta\theta$ we select those samples $i$ for which $\text{CDF}_{\text{i}}^{\text{random}}(\Delta\theta) \geq \text{CDF}^{\text{experiment}}$. If for a particular $\Delta\theta$ the total number of such selected samples is $m(\Delta\theta)$, the probability of observing equal or higher bias towards alignment of the two angles from a random experiment than that actually observed experimentally is $P(\Delta\theta) = m(\Delta\theta)/M$ (Figure 5 – figure supplement 1d). It can clearly be seen that the percentage probability of observing such high values of CDF for smaller $\Delta\theta < 30°$ from a completely random sampling of $\Delta\theta$ is $\approx 1$, i.e., approximately one sample of $N = 19$ experiments for every 100 samples, a very low number. The few colonies for which $\theta_{\text{nematic}}(0\text{ hr})$ does not match well with $\theta_{\text{colony}}(2\text{ hr})$ appear to align with the orientation $\theta_{\text{colony}}(0\text{ hr})$ of tissue initial elongation – the associated mechanism is, however, still open (see Figure 5 – figure supplement 1e). This entire analysis clearly indicates that the orientation of the mean cell-shape nematic at the initial time has correlation with the final anisotropy orientation of the colony with a high probability.

#### II. VERTEX MODEL SIMULATION

A planar vertex model is one of the most commonly used computational tool to understand mechanics of epithelial monolayers [13]. In this description, the tissue is assumed to be a 2-D sheet composed of cells which are represented by planar polygons sharing vertices and edges with their neighbors (Figure 7a). Governing equations are formulated to study the dynamics of evolution of each vertex of a cell. The total effective *energy* or work-function $W$ of the tissue arises from area deformation of the cells and a combination of acto-myosin contractility and membrane adhesion energy at the cell interfaces (Figure 7b).

$$W = \sum_{\alpha=1}^{N}\frac{K_\alpha}{2}(A_\alpha - A^0_\alpha)^2 + \sum_{<\alpha\beta>}\Lambda_{\alpha\beta}l_{\alpha\beta}, \tag{19}$$

where for a particular cell $\alpha$, $A_\alpha$ is the actual area and $A^0_\alpha$ is the preferred cell area. $l_{\alpha\beta}$ is the length of the bond connecting shared between the cells $\alpha$ and $\beta$ and $\Lambda_{\alpha\beta}$ is the effective contractility of that bond [14]. Note that the energy $W$ depends both on the position $\mathbf{r}_i$ of the vertices and cell connectivity.

For this model, the force $\mathbf{F}^i_{\text{vm}}$ acting on a particular vertex $i$ can be obtained by taking a derivative of $W$ [13, 15] with respect to the vertex position $\mathbf{r}_i$ as,

$$\mathbf{F}^i_{\text{vm}} = -\frac{\partial W}{\partial \mathbf{r}_i} \tag{20a}$$

$$= \eta \frac{d\mathbf{r}_i}{dt}, \tag{20b}$$

and is balanced by the friction force on the vertex from the substrate. Here, $\eta$ represents effective substrate viscosity experienced by the vertex via cell-substrate friction [15]. A mathematically equivalent description of the vertex model could also be made by using mechanical equilibrium or principle of virtual work [14, 16].

##### A. Inclusion of active cell stress in the vertex model formulation

Collectively migrating epithelial cells can have an active stress of the form [17]

$$\boldsymbol{\sigma} = \sigma_a(\hat{\mathbf{p}}\hat{\mathbf{p}} - \frac{1}{2}\mathbf{I}), \tag{21}$$

where for $\sigma_a > 0$ and $\sigma_a < 0$, respectively, for contractile and extensile cell active stress. When $\sigma_a > 0$, the cell has a tendency to contract along its polarisation $\hat{\mathbf{p}}$ and elongated perpendicular it, and vice-versa when $\sigma_a < 0$. The goal of this section is to present how this stress within a particular cell can be accommodated in the vertex model formalism by obtaining the corresponding active force $\mathbf{F}^i$ on the vertex $i$ of the cell (Figure 7c).

###### *1. Formulation of obtaining virtual strain of the cell from virtual displacement of cell vertices*

We note that, when the vertices are provided virtual displacement of $\{\delta x^i_\alpha\}$, the total internal virtual work because of the action of the cell active stress $\boldsymbol{\sigma}^\alpha$ on the resulting virtual deformation gradient $\delta\mathbf{u}$ is

$$\delta W_{\text{active}} = A\sigma_{\alpha\beta}\delta u_{\alpha\beta} = -\sum_i F^i_\alpha \delta x^i_\beta, \tag{22}$$

where $A$ is the instantaneous area of the cell. We note that the negative sign in front of the summation sign arises since $F^i_\alpha$ is taken as the force exerted by the interior of the cell on the vertices.

If we can find a relation between $\delta u_{\alpha\beta}$ and $\{\delta x^i_\alpha\}$, then Eq. 22 will provide us with expression for $F^i_\alpha$. We first note that, assuming an affine virtual displacement field, the virtual displacement $\delta x^i_\alpha$ of any vertex $i$ of a given cell $k$ will be given as

$$\begin{aligned} \delta x^i_\alpha &= \delta u_{\alpha\beta}(x^i_\beta - x^c_\beta) + \delta x^c_\alpha, \text{ and} \\ \delta \rho^i_\alpha &= \delta u_{\alpha\beta}\rho^i_\beta, \end{aligned} \tag{23}$$

where the superscript $c$ denotes the centroid of vertices, with position

$$x^c_\alpha = \frac{1}{N_{\text{ver}}}\sum_{i\in\text{cell}} x^i_\alpha, \tag{24}$$

the virtual displacement of this centroid is

$$\delta x^c_\alpha = \frac{1}{N_{\text{ver}}}\sum_{i\in\text{cell}} \delta x^i_\alpha, \tag{25}$$

and

$$\begin{aligned} \delta\rho^i_\alpha &= \delta x^i_\alpha - \delta x^c_\alpha, \\ \rho^i_\alpha &= x^i_\alpha - x^c_\alpha. \end{aligned} \tag{26}$$

Here, $N_{\text{ver}}$ is the number of vertices of the cell under consideration. The connection between $\delta\rho^i_\alpha$ and $\delta u_{\alpha\beta}$ as per Eq. 23 provides a set of $2 \times (N_{\text{ver}} - 1)$, independent linear equations, and can be guaranteed a unique solution in terms of the four independent components of the tensor $\delta u_{\alpha\beta}$, i.e., $\delta u_{11}, \delta u_{12}, \delta u_{21}$ and $\delta u_{22}$ only for $N_{\text{ver}} = 3$. For $N_{\text{ver}} > 3$, we obtain a solution in the sense of least squares. To do that we minimise the error

$$S = \sum_{i \in \text{cell}} (\delta\rho^i_\alpha - \delta u_{\alpha\beta}\rho^i_\beta)(\delta\rho^i_\alpha - \delta u_{\alpha\gamma}\rho^i_\gamma), \tag{27}$$

where repeated summation is implied over the Greek numerals $\alpha$, $\beta$ and $\gamma$, which can take values of 1 and 2. This error $S$ is to be minimised over $\delta u_{\mu\nu}$, and will give four independent linear equations. Differentiating $S$ with respect to $\delta u_{\mu\nu}$, we get the following set of equations (in indicial notation)

$$\frac{\partial S}{\partial \delta u_{\mu\nu}} = -2 \sum_{i \in \text{cell}} (\rho^i_\nu \delta\rho^i_\mu - \delta u_{\mu\beta}\rho^i_\beta \rho^i_\nu) = 0. \tag{28}$$

These equations can be written in more readable format in the following manner.

$$\begin{bmatrix} \delta\rho_{11} \\ \delta\rho_{21} \\ \delta\rho_{12} \\ \delta\rho_{22} \end{bmatrix} = \begin{bmatrix} \rho_{11} & \rho_{12} & 0 & 0 \\ \rho_{12} & \rho_{22} & 0 & 0 \\ 0 & 0 & \rho_{11} & \rho_{12} \\ 0 & 0 & \rho_{12} & \rho_{22} \end{bmatrix} \begin{bmatrix} \delta u_{11} \\ \delta u_{12} \\ \delta u_{21} \\ \delta u_{22} \end{bmatrix} \tag{29}$$

where the quantities are defined as follows:

$$\begin{aligned} \rho_\alpha &= \sum_i \rho^i_\alpha = 0, \\ \rho_{\alpha\beta} &= \sum_i \rho^i_\alpha \rho^i_\beta, \\ \delta\rho_{\alpha\beta} &= \sum_i \rho^i_\alpha \delta\rho^i_\beta, \end{aligned} \tag{30}$$

such that $\alpha \in \{1, 2\}$. Equation 29 can be solved to provide the components of $\delta U$ as follows.

$$\begin{aligned} \delta u_{11} &= \frac{\rho_{12}\delta\rho_{21} - \rho_{22}\delta\rho_{11}}{\rho_{12}^2 - \rho_{11}\rho_{22}}, \\ \delta u_{12} &= \frac{\rho_{12}\delta\rho_{11} - \rho_{11}\delta\rho_{21}}{\rho_{12}^2 - \rho_{11}\rho_{22}}, \\ \delta u_{21} &= \frac{\rho_{12}\delta\rho_{22} - \rho_{22}\delta\rho_{12}}{\rho_{12}^2 - \rho_{11}\rho_{22}}, \\ \delta u_{22} &= \frac{\rho_{12}\delta\rho_{12} - \rho_{11}\delta\rho_{22}}{\rho_{12}^2 - \rho_{11}\rho_{22}}. \end{aligned} \tag{31}$$

Thus in Eq. 31 we have the virtual deformation tensor for a given cell $k$ in terms of the virtual displacement of its vertices $\{\delta\rho^i_\alpha\}$ with respect to its vertex centroid. These expressions are to be substituted in Eq. 22 to find the forces that are exerted on cell vertices due to the active cell stress in Eq. 21.

##### 2. *Connection between cell active stress and vertex forces*

The active stress for the cell in Eq. 21 can be written in indicial notation as

$$\sigma_{\alpha\beta} = \sigma_a(p_\alpha p_\beta - \frac{1}{2}\delta_{\alpha\beta}), \tag{32}$$

and the contribution to the virtual work is

$$\delta W_{\text{active}} = A\sigma_{\alpha\beta}\delta u_{\alpha\beta}. \tag{33}$$

Using the expression for $\delta u_{\alpha\beta}$ from Eq. 31 we get

$$\begin{aligned} A\sigma_{\alpha\beta}\delta u_{\alpha\beta} &= A\sigma_{11}\delta u_{11} + A\sigma_{12}(\delta u_{12}+\delta u_{21}) + A\sigma_{22}\delta u_{22} \\ &= A\sigma_a(p_1^2-1/2)\left(\frac{\rho_{12}\delta\rho_{21}-\rho_{22}\delta\rho_{11}}{\rho_{12}^2-\rho_{11}\rho_{22}}\right) \\ &+ A\sigma_a(p_2^2-1/2)\left(\frac{\rho_{12}\delta\rho_{12}-\rho_{11}\delta\rho_{22}}{\rho_{12}^2-\rho_{11}\rho_{22}}\right) \\ &+ A\sigma_a(p_1p_2)\left(\frac{\rho_{12}\delta\rho_{11}-\rho_{11}\delta\rho_{21}}{\rho_{12}^2-\rho_{11}\rho_{22}} + \frac{\rho_{12}\delta\rho_{22}-\rho_{22}\delta\rho_{12}}{\rho_{12}^2-\rho_{11}\rho_{22}}\right), \\ &= -\sum_{\text{vertices } i} F_\alpha^i \delta x_\alpha^i. \end{aligned} \tag{34}$$

We now note from Eq. 30 that

$$\delta\rho_{\alpha\beta} = \sum_i \rho_\alpha^i(\delta x_\beta^i - \delta x_\beta^c) = \sum_i \rho_\alpha^i \delta x_\beta^i. \tag{35}$$

Using Eq. 35, Eq. 34 is rewritten as

$$\begin{aligned} A\sigma_{\alpha\beta}\delta u_{\alpha\beta} = A\sigma_a \sum_i \Bigg[ &\frac{(p_1^2-1/2)\left[\rho_{12}\rho_2^i-\rho_{22}\rho_1^i\right] + (p_1p_2)\left[\rho_{12}\rho_1^i-\rho_{11}\rho_2^i\right]}{\rho_{12}^2-\rho_{11}\rho_{22}}\delta x_1^i \\ &+ \frac{(p_2^2-1/2)\left[\rho_{12}\rho_1^i-\rho_{11}\rho_2^i\right] + (p_1p_2)\left[\rho_{12}\rho_2^i-\rho_{22}\rho_1^i\right]}{\rho_{12}^2-\rho_{11}\rho_{22}}\delta x_2^i \Bigg]. \end{aligned} \tag{36}$$

Hence, from Eqs. 36 and 34 the contribution to the force on vertex $i$ due to the cell active stress from the cell is

$$\begin{aligned} F_1^i &= -A\sigma_a \frac{(p_1^2-1/2)\left[\rho_{12}\rho_2^i-\rho_{22}\rho_1^i\right] + (p_1p_2)\left[\rho_{12}\rho_1^i-\rho_{11}\rho_2^i\right]}{\rho_{12}^2-\rho_{11}\rho_{22}}, \\ F_2^i &= -A\sigma_a \frac{(p_2^2-1/2)\left[\rho_{12}\rho_1^i-\rho_{11}\rho_2^i\right] + (p_1p_2)\left[\rho_{12}\rho_2^i-\rho_{22}\rho_1^i\right]}{\rho_{12}^2-\rho_{11}\rho_{22}}. \end{aligned} \tag{37}$$

It can very easily be seen from Eq. 37 that for a given cell, the sum of all the active forces on the vertices of that cell will be zero. Moreover, the total torque of all these forces with respect to the vertex centroid as defined in Eq. 24 can also be shown to be equal to zero. This is as expected from a symmetric stress tensor. In the current work, we take the cell active stress as extensile, and hence $\sigma_a < 0$ [9].

Finally, the contribution of this active force to any vertex is a vector sum of the force contributions coming from all the cells that the vertex is a part of, and is given by,

$$(F_\alpha^i)^{\text{total}} = \sum_{\text{cell } k\in\text{vertex } i} (F_\alpha^i)^{\text{total}}; \quad \alpha \in \{1,2\}. \tag{38}$$

The active force on the vertex would be added to the forces from cell area deformation and cell edge contractility as discussed in the earlier section.

##### B. Polarised or biased edge tensions

It is observed that T1 transitions can be influenced by the polarity of the cells [18]. Such bias in T1 transitions can be introduced in the vertex model by modifying the tension $\lambda_{\alpha\beta}$ of any edge $\alpha\beta$ based on its relative orientation $\hat{\mathbf{e}}_{\alpha\beta}$ with respect to the polarity $\hat{\mathbf{p}}_\alpha$ and $\hat{\mathbf{p}}_\beta$ of the shared cells (Figure 7e) as

$$\Lambda_{\alpha\beta} = \Lambda_{\alpha\beta}^0 + \lambda(1 - [(\hat{\mathbf{e}}_{\alpha\beta}\cdot\hat{\mathbf{p}}_\alpha)^2 + \hat{\mathbf{e}}_{\alpha\beta}\cdot\hat{\mathbf{p}}_\beta)^2]) + \xi(t), \tag{39}$$

where the quantities $\Lambda_{\alpha\beta}^0$, $\lambda$, $\xi(t)$ are, respectively, the base edge tension, angle dependent edge tension bias, and correlated noise that is calculated as

$$\frac{d\xi(t)}{dt} = -\frac{1}{\tau}\xi(t) + \gamma(t), \text{ such that} \tag{40}$$

$$\langle\gamma(t)\rangle = 0, \text{ and} \tag{41}$$

$$\langle\gamma(t')\gamma(t'+t)\rangle = \frac{2\Delta\Lambda^2}{\tau}\delta(t). \tag{42}$$

Here, $\tau$ and $\Delta\Lambda$ are the correlation time and strength, respectively, for the correlated noise. This noise promotes T1 transitions and introduces tissue fluidity in the vertex model [19]. The angle dependent bias ensure that the edges that are parallel (perpendicular) to the polarisation of the cells have a tension that is lower (higher) by $\lambda$ than its base value $\Lambda^0_{\alpha\beta}$. This bias enhances the propensity of tissue shear along cell polarisation due to T1 transitions.

##### C. Inclusion of cell motility

In addition to the forces described in the previous section, cells can also generate motility force [20, 21]. Following the idea of self-propelled cells [22], any cell $\alpha$ can also be modeled to have a speed $v_0$, such that in the absence of any other forces it moves with a velocity $v_0\hat{\mathbf{p}}_\alpha$, where $\hat{\mathbf{p}}_\alpha$ is the polarisation direction of the cell as described earlier (Figure 7d). A simplest representation of motile force on a particular vertex $i$ is

$$\mathbf{F}^i_{\text{motile}} = \eta v_0 \frac{1}{N_i} \sum_{\text{cell }\beta} \hat{\mathbf{p}}_\beta, \tag{43}$$

where $N_i$ is the total number of cells that contain the particular vertex $i$, and $\beta$ is the index of each of these cells [23]. The motile force, if included, will be in addition to the forces as described in the previous sections.

##### D. Dynamical Evolution

Dynamical evolution in these simulations involves updating the position of vertex $i$ as per the following equation

$$\eta \frac{d\mathbf{r}_i}{dt} = \mathbf{F}^i_{\text{basic}} + \mathbf{F}^i_{\text{active}} + \mathbf{F}^i_{\text{motile}}, \tag{44}$$

where the subscripts basic, active, and motile correspond to the contributions to the forces, respectively, from basic vertex model, active stress and biased edge tensions, and cell motility. The evolution of edge tensions is as discussed in the previous section. In addition to the evolution of the vertex positions, whenever the size of any edge becomes smaller than a critical size $\epsilon_{\text{cls}}$, a connectivity change, termed as T1 transition in the literature [14, 15], is implemented – the size of the new edge $\epsilon_{\text{opn}} > \epsilon_{\text{cls}}$ (Figure 7f).

The next step is to update the polarisation $\hat{\mathbf{p}}$ of the cells. In our description, the $\hat{\mathbf{p}}^\alpha$ for a given cell $\alpha$ is modeled to have a tendency to align with the orientation of its neighbours (ngb) without distinction between $\hat{\mathbf{p}}^{\text{ngb}}$ and $-\hat{\mathbf{p}}^{\text{ngb}}$. i.e., nematically. This is achieved with the following simple differential equation for $\hat{\mathbf{p}}_\alpha$ of any cell $\alpha$ in the interior:

$$\frac{d\hat{\mathbf{p}}_\alpha}{dt} = (\xi m_{\text{align}} + \xi_s m_{\text{stretch}} + \xi_{\text{rand}})\hat{\mathbf{p}}^\perp_\alpha, \tag{45}$$

where $\hat{\mathbf{p}}^\perp$ is such that

$$\hat{\mathbf{e}}_z = \hat{\mathbf{p}}_\alpha \times \hat{\mathbf{p}}^\perp_\alpha,$$

the $\hat{\mathbf{e}}_z$ is the vector normal to the plane of the paper.

$$m_{\text{align}} = \hat{\mathbf{e}}_z \cdot \left[ \frac{1}{N_\alpha} \sum_{\beta\in\text{ngb}} \text{sign}(\hat{\mathbf{p}}_\alpha \cdot \hat{\mathbf{p}}_\beta)(\hat{\mathbf{p}}_\alpha \times \hat{\mathbf{p}}_\beta) \right]$$

is the rotational torque on $\hat{\mathbf{p}}_\alpha$ that aligns it with respect to the mean orientation of its neighbors and $\xi$ is the alignment rate.

$$m_{\text{stretch}} = \hat{\mathbf{e}}_z \cdot [\text{sign}(\hat{\mathbf{e}}_x \cdot \hat{\mathbf{p}}_\alpha)(\hat{\mathbf{p}}_\alpha \times \hat{\mathbf{e}}_x)]\hat{\mathbf{p}}^\perp_\alpha$$

is the rotational torque that aligns $\hat{\mathbf{p}}_\alpha$ along the orientation $\hat{\mathbf{e}}_x$ of periodic stretch when applicable and $\xi_s$ is the corresponding alignment rate.

The quantity $\text{sign}(\hat{\mathbf{p}}_\alpha \cdot \hat{\mathbf{p}}_\beta)$ ensures that $\hat{\mathbf{p}}_\alpha$ nematically align with polarisation of the $N_\alpha$ neighbours in a mean sense. Similarly, the quantity $\text{sign}(\hat{\mathbf{p}}_\alpha \cdot \hat{\mathbf{p}}_\beta)$ is to ensure that $\hat{\mathbf{p}}_\alpha$ aligns nematically along the direction of stretch (when applicable). The orientational noise $\xi_{\text{rand}}$ is modeled as uncorrelated Gaussian noise of zero mean and strength $D_r$.

This evolution rule also automatically ensures that the $\hat{\mathbf{p}} \cdot \hat{\mathbf{p}} = 1$ for all the cells. For any cells at the boundary, the only difference is that the alignment torque is

$$m_{\text{align}} = \hat{\mathbf{e}}_z \cdot \text{sign}(\hat{\mathbf{p}}_\alpha \cdot \hat{\mathbf{t}})(\hat{\mathbf{p}}_\alpha \times \hat{\mathbf{t}}),$$

where

$$\hat{\mathbf{t}} = \frac{\mathbf{r}_{\text{next}} - \mathbf{r}_{\text{before}}}{|\mathbf{r}_{\text{next}} - \mathbf{r}_{\text{before}}|},$$

and $\mathbf{r}_{\text{next}}$ and $\mathbf{r}_{\text{before}}$ correspond to the position vectors of the two contiguous cells of cell-$\alpha$ at the boundary.

The initial condition for the positions $\{\mathbf{r}\}$ of the vertices and the polarisation $\{\hat{\mathbf{p}}\}$ of the cells is generated as follows. A confluent colony of 332 cells, a number similar to that observed experimentally, is initially confined within a circular region of radius $R_{\text{colony}} \approx 320$ (see Supplementary Material Section IE for clarification regarding the simulation parameters.) The cells are provided with an initial polarisation $\hat{\mathbf{p}} = \cos\phi_p \hat{\mathbf{e}}_1 + \sin\phi_p \hat{\mathbf{e}}_2$ such that the orientation $\phi_p(x, y)$ is

$$\phi_p(x, y) = \frac{1}{2}\left(\tan^{-1}\left[\frac{y - y_1}{x - x_1}\right] + \tan^{-1}\left[\frac{y - y_2}{x - x_2}\right]\right) + \frac{\pi}{2}, \tag{46}$$

where $(x_1, y_1)$ and $(x_2, y_2)$ is the location of two $+1/2$ defects, which are taken to be symmetrically separated by a distance of $1.6R_{\text{colony}}$ with respect to the center of the circle (also see Supplementary Section ID). An orientation shift of $\pi$ is randomly added to $\phi_p$ of individual cells such that the vector sum of $\hat{\mathbf{p}}$ over all the cells is zero. This step is irrelevant for the implementation of forces due to cell active stress and biased edge tensions due to their inherent nematic symmetry (Eqs. 21 and 39) but ensures that the total motile force from all the cells is zero. The cells are then provided with small active stress and evolved under confinement for some time using the above rules till $\{\mathbf{r}\}$ and $\{\mathbf{p}\}$ are almost equilibrated. This configuration is then used as the initial condition for all the numerical experiments, in which the confinement is removed to allow the colonies to elongate (see Movie 7). Different simulation conditions correspond to varying proportion of active cell stress, biased cell edge tension and cell motility. The results from the simulations are then analysed in a similar manner as for the experimental data (Figure 7 – figure supplement 1).

This set of polarity evolution rules for the interior and the boundary cells ensure that in the steady-state the orientation field $\phi_p$ satisfies in a coarse-grained sense, i.e., on longer length-scales, the Laplace equation $\nabla^2\phi_p = 0$. The nematic field for the interior cells tends to have a uniform orientation, whereas the nematic field for the boundary cells tries to align along the colony boundary. Because of this boundary alignment tendency of the border cells, at least for the parameters used in our simulations (Supplementary Material Section IE), the total topological charge of the polarisation field is mostly conserved in our simulations, i.e., remains equal to the total initial charge of $+1$ (for a relevant study in another context see Ref. [24].) Due to the action of active cell stress (Eq. 21) the cells have a tendency to elongate predominantly along $\hat{\mathbf{p}}$ thus creating a cell-shape nematic orientation field $\mathbf{q} = \cos\phi\hat{\mathbf{e}}_1 + \sin\phi\hat{\mathbf{e}}_2$ (also see Supplementary Material Sections IA-C.) Hence, the cell elongation anisotropy, as described earlier in Supplementary Material Sections IC and ID, can loosely be thought of to be a readout of the internal polarisation of the cells. However, since the actual cell shape results from the interplay between cell mechanical properties and active cell stress, $\hat{\mathbf{p}}$ and $\mathbf{q}$ are not necessarily always fully aligned. Thus, for the cell-shape nematic field, the total topological charge can be less than $+1$ if the aligning tendency of internal cells dominates, e.g., when the cell active stress is large (see Movie 7).

##### E. Parameters used and the non-dimensional groups present in the Vertex Model

Using reference scales for length ($l_0$), time ($t_0$) and energy ($U_0$), we get the following set of non-dimensional groups for the various parameters that are involved in the simulations. The non-dimensional counterparts of the parameters used in the simulations have been assigned an additional $'$ to make them distinct from the original.

1. Substrate friction: $\frac{\eta l_0^2}{t_0 U_0} \to \eta' = 1000$
2. Area modulus: $\frac{K l_0^4}{U_0} \to K' = 0.1$
3. Cell preferred area: $\frac{A_0}{l_0^2} \to A_0' = 1000$
4. Edge contractility: $\frac{\Lambda_0 l_0}{U_0} \to \Lambda_0' = 100$, $\frac{\Delta\Lambda l_0}{U_0} \to \Delta\Lambda = 20$, $\frac{\lambda l_0}{U_0} \to \lambda' \in [0, 50]$

5. Cellular active stress (negative sign indicates extensile): $\frac{\sigma_a l_0^2}{U_0} \rightarrow \sigma'_a \in [-8, 4]$.
6. Motility: $\frac{v_0 t_0}{l_0} \rightarrow v'_0 = 0.05$
7. Polarisation alignment rate: $\xi t_0 \rightarrow \xi' = 0.03$
8. Stretch alignment rate: $\xi_s t_0 \rightarrow \xi' = 0.001$
9. Strength of Gaussian noise for polarisation: $t_0 D_r \rightarrow D'_r = 2.7 \times 10^{-6}$ (very small)
10. Relaxation time for edge tension fluctuations: $\frac{\tau}{t_0} \rightarrow \tau' = 10^3$
11. Total experimental time: $\frac{T_{\text{total}}}{t_0} \rightarrow T'_{\text{total}} = 4000$
12. T1 transition cutoffs: $\frac{\epsilon_{\text{opn}}}{l_0} \rightarrow \epsilon'_{\text{opn}} = 6$ and $\frac{\epsilon_{\text{cls}}}{l_0} \rightarrow \epsilon'_{\text{cls}} = 4$
13. Colony size: $\frac{R_{\text{colony}}}{l_0} \rightarrow R'_{\text{colony}} = 317$
14. Simulation time-step: $\frac{\Delta t}{t_0} \rightarrow \Delta t' = 1$

We can now choose the scales $l_0$, $t_0$, and $U_0$. We use the simulation time-step $\Delta t' = 1$. We get the values of total cumulative shear in the range $\approx 0.05 - 0.2$ that is comparable with the experimental values at the end of the 4000 simulation time. Since the total duration of the experiment is 2 hrs, this implies that $t_0 \approx 1.8$ s in experimental units. We also know from the experiments that the area of one cell is $\approx 400\ \mu\text{m}^2$. Since, within our simulations, the preferred area of cells is taken as $A_0 = 1000$, this implies that $l_0 \approx 0.63\ \mu$m.

The energy (or force) scale can be chosen arbitrarily since it is present in both the left hand and the right hand sides of Eq. 44, and must only be chosen consistently. We now figure out the relative contributions to the forces that can come from different components for a cell: motile force, active force, basic edge contractility, active edge contractility, and isotropic force. For the values used the non-dimensional (simulation) values are:

1. cell motile force: $\approx \eta' v'_0 = 1000 \times [0, 0.05] = [0, 50]$
2. cell active force (extensile): $\approx |\sigma'_a \sqrt{A'_0}| = [0, 8] \times \sqrt{1000} \approx [0, 250]$
3. basic edge tension: $\Lambda'_0 = 100$
4. biased edge tension: $\approx \lambda' = [0, 45]$
5. cell area stress: $K' \sqrt{A'_0} \Delta A' \approx 3.16 \times [0, 15] \approx [0, 45]$

One can see that forces from all these sources are of the same order of magnitude. Specifically when we wish to compare the relative magnitudes of active edge contractility and the active stress, a quantity of relevance is $\frac{\lambda'}{\sigma'_a \sqrt{A'_0}}$. The values of the active terms $\sigma'_a$, $\lambda'$ and $v'_0$ used in the simulations are shown in Figure 7 – figure supplement 1.


[1] B. Aigouy, R. Farhadifar, D. B. Staple, A. Sagner, J.-C. Röper, F. Jülicher, and S. Eaton, Cell **142**, 773 (2010).
[2] B. Aigouy, D. Umetsu, and S. Eaton, in *Drosophila* (Springer, 2016), pp. 227–239.
[3] M. Merkel, R. Etournay, M. Popović, G. Salbreux, S. Eaton, and F. Jülicher, Physical Review E **95**, 032401 (2017).
[4] R. Etournay, M. Merkel, M. Popović, H. Brandl, N. A. Dye, B. Aigouy, G. Salbreux, S. Eaton, and F. Jülicher, eLife **5**, e14334 (2016).
[5] E. Paluch and C.-P. Heisenberg, Curr. Biol. **19**, R790 (2009).
[6] M. Rauzi and P.-F. Lenne, Curr Top Dev Biol **95**, 93 (2011).
[7] R. Etournay, M. Popović, M. Merkel, A. Nandi, C. Blasse, B. Aigouy, H. Brandl, G. Myers, G. Salbreux, F. Jülicher, et al., eLife **4**, e07090 (2015).
[8] Z. Püspöki, M. Storath, D. Sage, and M. Unser, in *Focus on Bio-Image Informatics* (Springer, 2016), pp. 69–93.
[9] T. B. Saw, A. Doostmohammadi, V. Nier, L. Kocgozlu, S. Thampi, Y. Toyama, P. Marcq, C. T. Lim, J. M. Yeomans, and B. Ladoux, Nature **544**, 212 (2017).
[10] MATLAB, *version 9.4.0 (R2018a)* (The MathWorks Inc., Natick, Massachusetts, 2018).
[11] M. Kleman and O. D. Laverntovich, *Soft matter physics: an introduction* (Springer Science & Business Media, 2003).
[12] P.-G. De Gennes and J. Prost, *The physics of liquid crystals*, vol. 83 (Oxford university press, 1995).
[13] R. Farhadifar, J.-C. Röper, B. Aigouy, S. Eaton, and F. Jülicher, Curr. Biol. **17**, 2095 (2007).

[14] S. Alt, P. Ganguly, and G. Salbreux, Philosophical Transactions of the Royal Society B: Biological Sciences **372**, 20150520 (2017).
[15] A. G. Fletcher, M. Osterfield, R. E. Baker, and S. Y. Shvartsman, Biophys. J. **106**, 2291 (2014).
[16] H. H. Chen and G. W. Brodland, Journal of biomechanical engineering **122**, 394 (2000).
[17] M. Popović, A. Nandi, M. Merkel, R. Etournay, S. Eaton, F. Jülicher, and G. Salbreux, New Journal of Physics **19**, 033006 (2017).
[18] K. Sato, T. Hiraiwa, E. Maekawa, A. Isomura, T. Shibata, and E. Kuranaga, Nature communications **6**, 10074 (2015).
[19] S. Curran, C. Strandkvist, J. Bathmann, M. de Gennes, A. Kabla, G. Salbreux, and B. Baum, bioRxiv p. 078204 (2016).
[20] A. Mogilner and G. Oster, Biophys. J. **71**, 3030 (1996).
[21] T. D. Pollard and G. G. Borisy, Cell **112**, 453 (2003).
[22] D. Bi, X. Yang, M. C. Marchetti, and M. L. Manning, Physical Review X **6**, 021011 (2016).
[23] D. M. Sussman, Computer Physics Communications **219**, 400 (2017).
[24] R. M. W. van Bijnen, R. H. J. Otten, and P. van der Schoot, Phys. Rev. E **86**, 051703 (2012).

**Video 1**. **Spontaneous symmetry breaking of circular colonies**. Time-lapse of a MDCK cells colony freely evolving after removal of a PDMS stencil. Time in hh:mm. Scale bar 50 μm.

**Video 2**. **Cyclic stretching of MDCK colonies**. Time-lapse of a MDCK colony under cyclic stretching (5% and 120s). Time in hh:mm:ss. Scale bar 50 μm.

**Video 3**. **Expansion of MDCK colonies under cyclic stretching**. Time-lapse of a MDCK colony under cyclic stretching (5% and 60s) at the 0% strain position. Time in hh:mm. Scale bar 50 μm.

**Video 4**. **Early symmetry breaking of circular colonies**. Composite array time-lapse of MDCK-GFP-Ecadherin cells colonies during the first two hours of expansion. Time in hh:mm. Scale bar 50 μm.

**Video 5**. **Disruption of the acto-myosin cable**. Movie showing the disruption of the acto-myosin cable. The colony and the micropipette used are shown at the left and the myosin signal is shown at the right. Cytochalasin D was mixed with cy5 to allow its visualization. Time in hh:mm. Scale bar 50 μm (right).

**Video 6**. **Nematic field alignment precedes colony elongation**. Movie showing cell-shape nematics and topological defects of an elongating colony. Time in hh:mm. Scale bar 50 μm.

**Video 7**. **Contributions to colony elongation**. Movie showing cell tracking, neighbor triangulation, cell-shape nematics and topological defects of an elongating colony. Time in hh:mm. Scale bar 50 μm.

**Video 8**. ***In silico* recreation of a colony elongation**. Vertex model simulation of a colony elongation. Cell-shape nematics, topological defects and cell polarization are followed over time.

**Video 9. *In silico* recreation of single versus collective effect of stretching.** Competition between active cell stress $\boldsymbol{\sigma}_a$ and biased junction tension $\lambda$ governs direction of colony elongation. (**Left**) When cell-cell junctions are normal, we propose that $\lambda'$ = 50 dominates over $\boldsymbol{\sigma}_a'$ = -2 and the colony elongates along the direction of stretch (horizontal) collectively through T1 transitions (**collective** stretching). (**Right**) When E-cadherin levels are low (blocked by anti-E-cadherin antibody), the effect of edge tensions $\lambda'$ = 25 is lowered as compared and that of active cell stress $\boldsymbol{\sigma}_a'$ = -4 is increased, thus leading to elongation of the colony perpendicular to the direction of stretch (vertical) through individual cell elongation (**single** stretching). The superscript ’ indicates non-dimensionalised parameter (see Supplementary Information Section IE).
